# Glutathionylation of Pyruvate Dehydrogenase Complex E2 and Inflammatory Cytokine Production During Acute Inflammation Are Magnified By Mitochondrial Oxidative Stress

**DOI:** 10.1101/2023.01.26.525791

**Authors:** David L. Long, Charles E. McCall, Leslie B. Poole

**Author notes:** Co-Corresponding Authors: Leslie B. Poole, Contact, Charles E. McCall.

## Abstract

Lipopolysaccharide (LPS) is a known inducer of inflammatory signaling which triggers generation of reactive oxygen species (ROS) and cell death in responsive cells like THP-1 promonocytes and freshly isolated human monocytes. A key LPS-responsive metabolic pivot point is the 9 megadalton mitochondrial pyruvate dehydrogenase complex (PDC), which provides pyruvate dehydrogenase (E1), lipoamide-linked transacetylase (E2) and lipoamide dehydrogenase (E3) activities to produce acetyl-CoA from pyruvate. While phosphorylation-dependent decreases in PDC activity following LPS treatment or sepsis have been deeply investigated, redox-linked processes have received less attention. Data presented here demonstrate that LPS-induced reversible oxidation within PDC occurs in PDCE2 in both THP-1 cells and primary human monocytes. Knockout of PDCE2 by CRISPR and expression of FLAG-tagged PDCE2 in THP-1 cells demonstrated that LPS-induced glutathionylation is associated with wild type PDCE2 but not mutant protein lacking the lipoamide-linking lysine residues. Moreover, the mitochondrially-targeted electrophile MitoCDNB, which impairs both glutathione-and thioredoxin-based reductase systems, elevates ROS similar to LPS but does not cause PDCE2 glutathionylation. However, LPS and MitoCDNB together are highly synergistic for PDCE2 glutathionylation, ROS production, and cell death. Surprisingly, the two treatments together had differential effects on cytokine production; pro-inflammatory IL-1β production was enhanced by the co-treatment, while IL-10, an important anti-inflammatory cytokine, dropped precipitously compared to LPS treatment alone. This new information may expand opportunities to understand and modulate PDC redox status and activity and improve the outcomes of pathological inflammation.

**Highlights:**

- PDCE2 is glutathionylated (-SSG) during acute inflammation in monocytes
- Lipopolysaccharide-induced PDCE2-SSG occurs in THP1 cells and human monocytes.
- Lipoamide-deficient PDCE2 lowers LPS-induced PDCE2-SSG and ROS production.
- MitoCDNB leads to ROS production without PDCE2-SSG, but is synergistic with LPS
- MitoCDNB causes the LPS-stimulated cytokine profile to become more proinflammatory

**Graphical Abstract:** 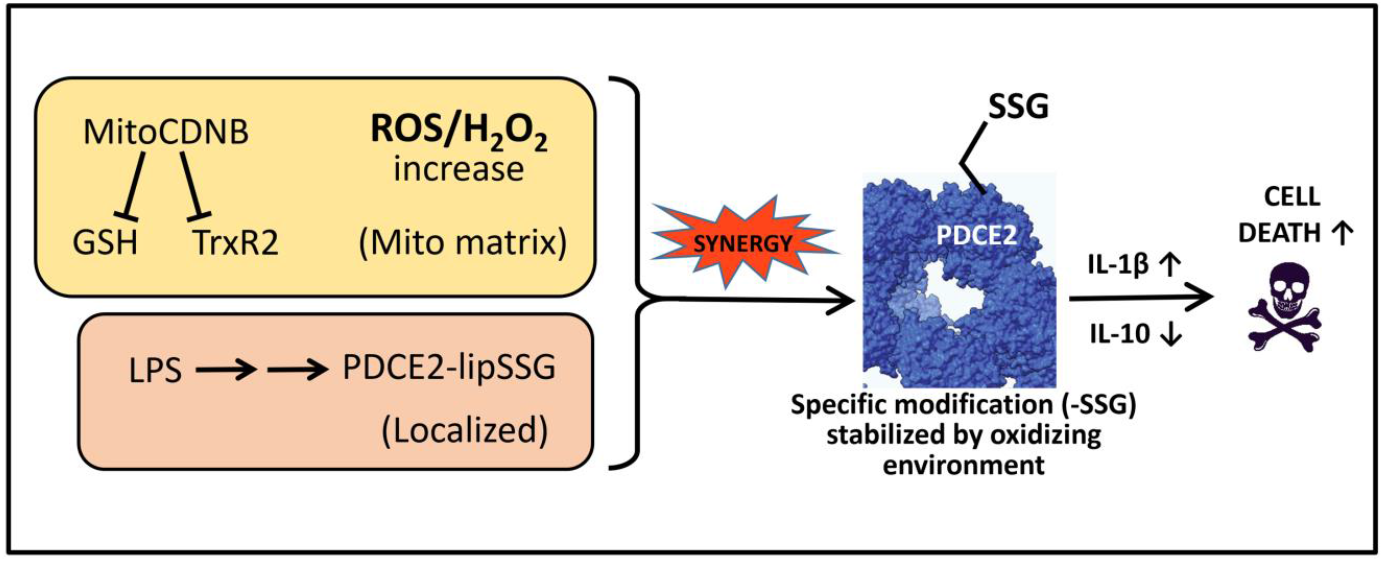

## 1. INTRODUCTION

Mitochondrially-produced reactive oxygen species (ROS)^1^ are important for cellular regulation in many scenarios, but are especially critical in the regulation of immune function including toll-like receptor (TLR) signaling and inflammatory responses [1, 2]. Known sources of mitochondrial ROS production include electron transport chain (ETC) components and NADPH oxidase activation [3]; however, it has also been established that 2-oxo acid dehydrogenases, particularly pyruvate dehydrogenase complex (PDC) and 2-oxoglutarate dehydrogenase complex (OGDHC, also known as α–ketoglutarate dehydrogenase), are oxidant generators under permissive conditions and are themselves subject to redox modifications [4-9]. Mailloux et al. recently reported that glutathionylation of PDC *in vitro* enhances its ROS-generating activity; a similar effect was also observed with OGDH glutathionylation [10]. Such redox modifications have important metabolic consequences given the key roles these enzymes play in aerobic glucose metabolism.

PDC and OGDHC are multicomponent enzymes that form large supramolecular structures and catalyze the oxidative decarboxylation of substrates linked to NAD^+^ reduction in mitochondria [4, 11]. Under aerobic conditions, the primary activities of PDC and OGDHC enzyme complexes become key metabolic regulatory nodes as they control: (i) for PDC, the conversion of pyruvate to acetyl-CoA, a key TCA carbon source (Fig. 1A), or (ii) for OGDHC, the conversion of 2-oxoglutarate to succinyl-CoA in the TCA cycle. NADH from both processes drives ATP production through the ETC. PDC, of interest here, is composed of three catalytic components, with the E2 component (along with E3 binding protein) forming the structural core. E1, made up of α and β subunits, catalyzes the thiamine diphosphate-dependent decarboxylation of pyruvate to release CO_2_ and form a high-energy, 2-carbon intermediate that is transferred to oxidized E2 lipoyl groups. Subsequent transfer of the acetyl group to coenzyme A (CoA) forms the central metabolite acetyl-CoA. Finally, the NAD^+^-dependent lipoamide dehydrogenase activity of FAD-containing E3 returns the lipoyl sulfurs to their oxidized, disulfide-bonded state for another round of catalysis. E3 is also the site where, under permissive conditions, NADH-driven oxidase activity produces superoxide and hydrogen peroxide from molecular oxygen (Fig. 1B and Fig. S1). While regulatory phosphorylation of PDC has been well established in sepsis, even leading to a promising new therapeutic approach [12-17], much remains to be learned about redox regulation of PDC.

**Figure 1.**
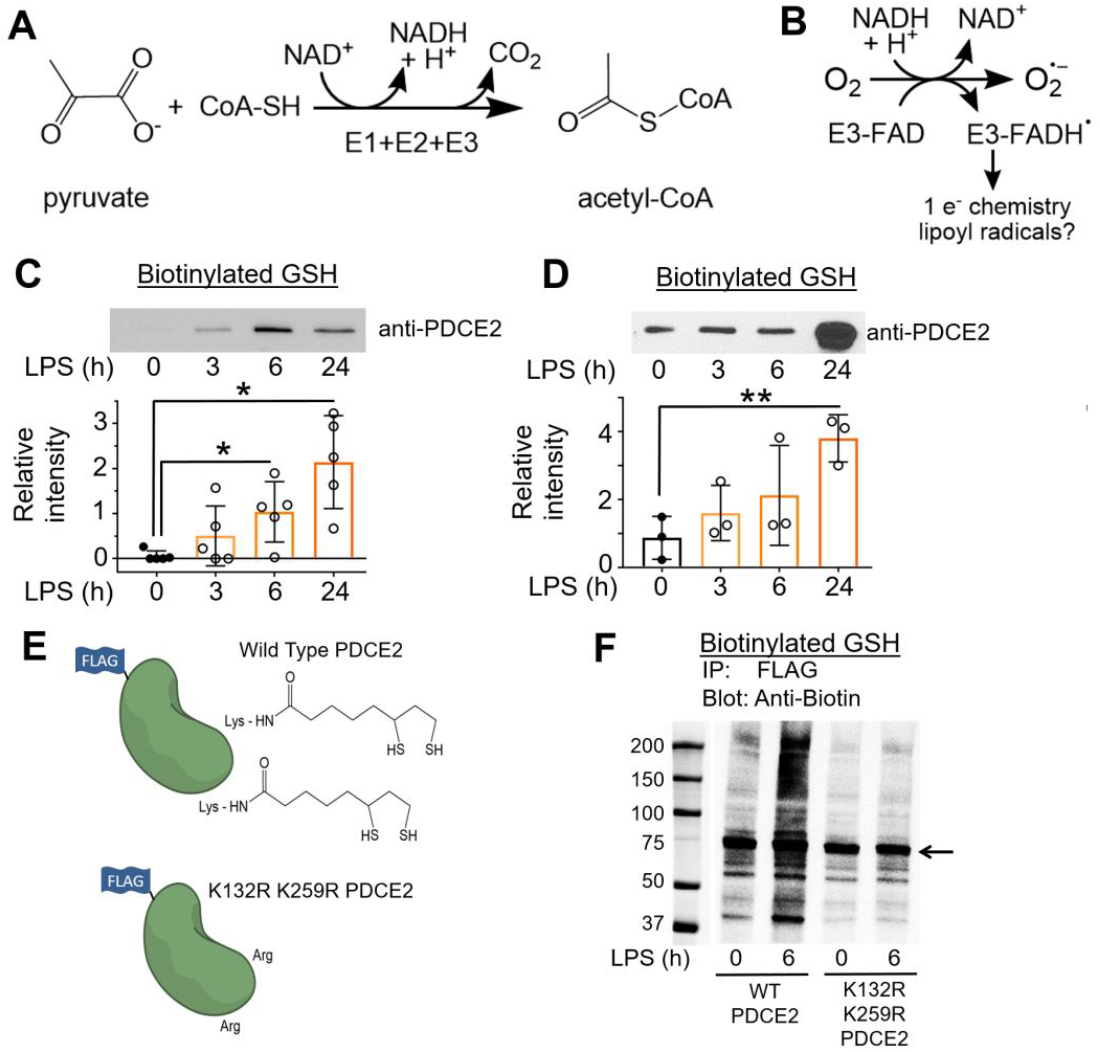
Functions of pyruvate dehydrogenase complex (PDC) proteins, and glutathionylation of monocyte PDCE2 in response to LPS. **(A)** Three components (E1, E2 and E3) of PDC together catalyze the decarboxylation of pyruvate to produce acetyl-CoA. **(B)** PDCE3 under permissive conditions exhibits NADH-dependent oxidase activity, producing superoxide and additional radical interactions. **(C)** Biotinylated glutathione (GSH) indicated glutathionylation of PDCE2 occuring in THP-1 pro-monocytes after treatment with 1 μg/mL LPS. After affinity capture using streptavidin beads and elution with dithiothreitol, PDCE2 was visualized by immunoblotting (n=5). A significant increase in glutathionylated PDCE2 occurred at 6 h and 24 h LPS treatment (* p<0.05). **(D)** Human primary monocytes treated with 100 ng/mL LPS (n=3) significantly increased glutathionylated PDCE2 at 24 h (** p<0.01). **(E)** A PDCE2 expression plasmid including sequences encoding an N-terminal FLAG tag was designed to replace lipoylated lysine residues (K132 and K259) with arginine residues, enabling expression of wild type (WT) and non-lipoylated PDCE2. **(F)** Using CRISPR-generated PDCE2 knockout THP-1 cells as host, plasmid-transfected cells expressing WT or K132R/K259R PDCE2 were treated with LPS or buffer for 6 h, followed by immunoprecipitation (IP) of the lysates for the FLAG-tagged proteins and immunoblotting for biotin to detect glutathionylation (Bio-GEE incorporation) (n=5).

Keenly aware that redox regulation of PDC in the face of the changing redox landscape of sepsis needs to be better understood, we set out to evaluate the glutathionylation status of PDC as it changes during acute inflammation stimulated by lipopolysaccharide (LPS) treatment (a model of sepsis) in human THP-1 promonocytic cells in culture and in freshly isolated monocytes from human blood. Glutathionylation associated with PDCE2 was indeed observed upon LPS treatment in both cases, but was compromised in recombinant expression systems with LPS-treated THP-1 cells where the lipoamide-linking lysine residues in PDCE2 were mutated. Treatment with the mitochondrially-targeted electrophile MitoCDNB alone, which compromised both glutathione-and thioredoxin-dependent reductase systems, did not lead to PDCE2 glutathionylation. However, its coadministration with LPS greatly augmented PDCE2 glutathionylation and associated cell death, suggesting cross-talk between reductase systems and PDC redox regulation. Moreover, cotreatment with LPS and MitoCDNB caused THP-1 cells to increase production of pro-inflammatory cytokine IL-1β but strongly suppress anti-inflammatory cytokine IL-10. This new information may expand opportunities to understand and modulate PDC redox status and activity and improve sepsis outcomes.

## 2. MATERIALS AND METHODS

See Supplementary Methods and Data for details.

### 2.1. Human Promonocytic THP-1 Cell Model and Isolation of Human Primary Monocytes

THP-1 cells were maintained in complete RPMI 1640 medium including 10% fetal bovine serum (FBS) and treated with 1 μg/mL bacterial lipopolysaccharide (LPS) as a model of acute inflammation and sepsis. Primary lymphocytes from heparinized venous blood were obtained by removal of RBCs, platelets, and neutrophils through Isolymph centrifugation, then monocytes were isolated by removing nonadherent cells after 2 h. Cells after 18 h incubation in fresh media were challenged with 100 ng/mL LPS.

### 2.2. Plasmid Constructs, Transfection and CRISPR Knockout Cells

A PDCE2 expression plasmid from SinoBiological (HG15002-UT) served as template for insertion of an N-terminal FLAG tag sequence and for generating a double mutant substituting Arg for two Lys residues (at 132 and 259). THP-1 cells were transfected with plasmid using GeneX Plus transfection reagent. PDCE2 CRISPR knockout THP-1 cells were commercially produced (Synthego Corporation).

### 2.3. Detection of Glutathionylated Proteins

Cells were preincubated for 1 h with culture medium containing 250 μM biotin-glutathione ethyl ester (BioGEE, ThermoFisher), then treated as indicated; they were then washed with ice-cold PBS and lysed in a RIPA buffer containing 10 mM *N*-ethylmaleimide (NEM) to block free thiols. NEM was removed with a desalting column, protein concentration was determined, and equivalent amounts of lysate proteins were incubated for 1 h at 4°C with streptavidin-conjugated magnetic beads. Beads were washed stringently (with 1M NaCl, 2M urea and 0.1% SDS, see Supplement), then *S*-glutathionylated proteins were released and prepared for SDS polyacrylamide gel electrophoresis (SDS-PAGE) with reducing sample buffer at 100°C (ThermoFisher #39000); immunoblotting was performed with anti-PDCE2 antibodies (GeneTex #GTX109766). Prestained Protein Standards (BioRad Precision Plus Protein™ Kaleidoscope™ Standards #1610375) were included on the SDS gels. In some experiments, mitochondrially-targeted 1-chloro-2,4-dinitrobenzene (MitoCDNB) was added at 10 μM along with BioGEE. For experiments involving plasmids, cells were transfected 36 h prior to the BioGEE incubation, then after treatment and lysis, samples were immunoprecipitated with anti-FLAG magnetic beads for 1 h, washed three times with lysis buffer, then eluted with low pH elution buffer. Samples were prepared with reducing or non-reducing sample buffer, then resolved by SDS-PAGE and immunoblotted for biotin. Note that while these approaches allow sensitive detection of stimulus-induced glutathionylation, the identification of the modified proteins is not direct and depends in part on stringent washing of the beads prior to elution of the biotinylated protein (linked via disulfide bond) by reductant.

### 2.4. ROS Assays

#### MitoSox Red ROS indicator

For most assays (endpoint assays), cells were treated for the specified time, then collected and counted, normalizing to 2×10^6^ cells per sample. They were washed once with Phenol Red Free media, incubated with 5 μM MitoSox Red for 20 min, washed, resuspended in Phenol Red Free media and plated in a black 96-well plate; fluorescence was measured with λ_ex_ and λ_em_ at 510nm and 580nm, respectively. In some assays, 10 μM MitoCDNB was added with the MitoSox prior to LPS treatment. For assays monitoring ROS production over time, cells were pre-incubated with MitoSox, then treated and monitored over 6 h at 30 min intervals.

#### Amplex Red H_2_O_2_ indicator

Cells were washed with Phenol Red Free media and resuspended at 1×10^6^/mL density into media containing Amplex Red (supplemented with horseradish peroxidase) at a final concentration of 100 μM, then plated as above and stimulated with LPS; absorbance was measured at 560 nm 6 h later.

### 2.5 Cell Death Assays

#### Fluorescent Live Cell/Dead Cell Assay

Cells were treated and plated as above for MitoSox endpoint assays, incubated with Multiplex Fluorometric Cytotoxicity Assay reagent (Promega) for 45 min, then assessed with λ_ex_=400nm/λ_em_=505nm for live cells, and λ_ex_=485nm/λ_em_=520nm for dead cells.

#### Lactate Dehydrogenase (LDH) Release Assay

Homogenous Membrane Integrity Assay (Promega) was used to measure LDH released into the media by 5 × 10^5^ cells per well after 18 h incubation with LPS, with Cyto-Tox One fluorescence measured at λ_ex_=560nm/λ_em_=590nm.

### 2.6. Cytokine Assays

For determination of cytokine (IL-1β and IL-10) levels in the media, cells with or without 30 min pretreatment with 10 μM MCDNB and/or addition of 1 μg/mL LPS for 6 or 24 h were removed by centrifugation, and media was tested by ELISA according to the manufacturer’s instructions (BD OptEIA Human IL-10 ELISA set #3555157 and BD OptEIA Human IL-1B ELISA set 2 #557953).

### 2.7 Imaging and Statistical analysis

ImageJ software (NIH) was used for densitometry analysis of immunoblots; density values were corrected for variations in total protein loading using β-actin as control. Individual data points represent averages across duplicates or triplicates of analyzed samples from wells of cells treated separately, with biological variation considered by conducting each set of experiments over at least 3 to 6 different days. Statistical analyses conducted with GraphPad Prism 7.05 used One Way ANOVA with Tukey’s post hoc multiple comparison with α=0.05, and the Shapiro-Wilk test for normality. All data are represented as mean + SEM.

## 3. RESULTS AND DISCUSSION

### 3.1. PDCE2 thiol oxidation during acute inflammation

#### 3.1.1 Monocyte PDCE2 is glutathionylated in response to LPS

Based on the recent study showing that PDCE2 could undergo S-glutathionylation when challenged chemically with oxidizing agents like oxidized glutathione (GSSG) [10], we evaluated S-glutathionylation of PDCE2 in our sepsis model systems. For sensitive detection of glutathionylation, human primary monocytes or THP-1 cells were pretreated with the biotin-tagged glutathione analog BioGEE for 1 h, then LPS was added for various times and cells were harvested and lysed in the presence of the thiol blocker NEM, followed by affinity capture with streptavidin, stringent washing of beads, then elution of disulfide-bound proteins with DTT. Immunoblotting for PDCE2 revealed that LPS stimulation led to a significant increase in PDCE2 glutathionylation at 6 and 24 h after 1 μg/mL LPS addition for THP-1 cells (Fig. 1C), and for isolated human monocytes at 24 h after 100 ng/mL LPS addition (Fig. 1D).

#### 3.1.2 Mutation of PDCE2 Lys residues to prevent lipoylation interferes with PDCE2 glutathionylation

To aid in assessment of the importance of PDCE2 lipoyl groups in the glutathionylation of this protein during LPS signaling, we obtained CRISPR-generated PDCE2 knockout (KO) THP-1 cells (Fig. S2) and designed a FLAG-tagged expression plasmid to express back either wild type (WT) PDCE2 or a mutant wherein the two Lys residues (K132 and K259) which serve as attachment sites for the lipoyl groups were mutated to Arg (Fig. 1E). As the lipoyl groups of PDCE2 play catalytic rather than structural roles and are within flexible “swinging arm” domains that extend beyond the core and E1 binding domains of PDCE2, we expect that the complex remains largely or fully intact, even though the mutated PDCE2 proteins lacking lipoyl groups are inactive [10, 11].

Repeating the BioGEE labeling experiment above, but then immunocapturing FLAG-tagged PDCE2 followed by blotting for biotinylated proteins, we found that not only PDCE2 but also other apparently associated proteins were glutathionylated in the system expressing the WT protein after 6 h LPS stimulation, whereas significantly less glutathionylation was detected in the mutant-expressing cells (Fig. 1F, see Fig. S3 for confirmation of PDCE2 pulldown).

### 3.2 Impact of CRISPR-generated knockout of PDCE2 in THP-1 cells, or mutation of lipoylation sites, on LPS-induced ROS production and cell death

#### 3.2.1 PDCE2 knockout lowers LPS-induced ROS production and cell death

Chemically-induced glutathionylation of PDCE2 has been associated with an increase in NADH-dependent ROS production by PDCE3 within the complex [10], prompting us to investigate both ROS generation and cell death imparted by LPS treatment linked to PDCE2 expression. MitoSox was used to detect mitochondrial LPS-induced ROS (Fig. 2A); this chemical probe is sensitive to oxidant-induced fluorescence increases caused by superoxide as well as other oxidants and is widely used in studies of ROS and mitochondria [18, 19]. Also used for these studies was Amplex Red to evaluate extracellular H_2_O_2_ as mitochondrially-produced H_2_O_2_, when upregulated, may also be detected extracellularly. The data indicated that with or without LPS treatment, the KO cell population released less H_2_O_2_ than WT cells. In addition, WT cells responded to LPS with detectably higher H_2_O_2_ release, whereas the KO cells did not (Fig. 2B). Taken together, these probes showed that ROS, and specifically H_2_O_2_, are significantly increased by treatment of THP-1 cells with LPS for 6 h (Fig. 2A and 2B). Effects similar to those with MitoSox were also observed with MitoPY1 which selectively detects H_2_O_2_ produced inside mitochondria (Fig. S4).

**Figure 2.**
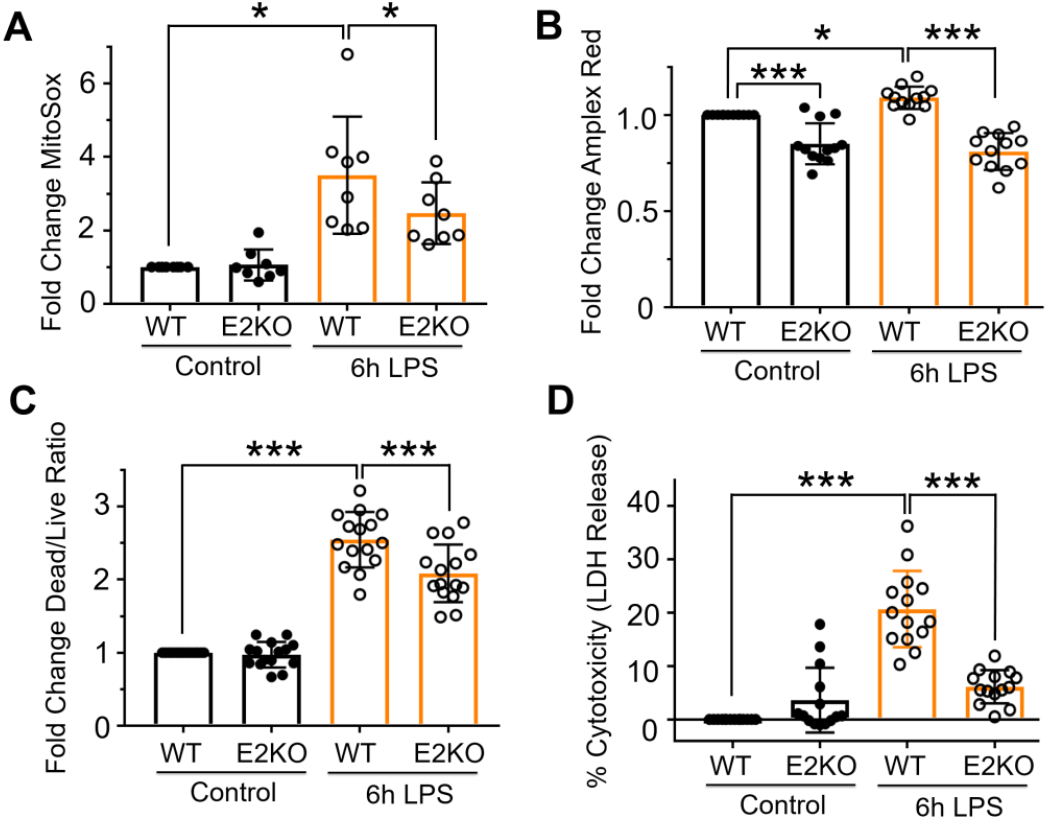
PDCE2 knockout lowers LPS-induced ROS production and cell death. (A) Mitochondrial ROS production was assessed in LPS-treated wild type (WT) THP-1 cells in comparison with CRISPR-mediated PDCE2 knockout (KO) THP-1 cells using 5 μM MitoSox Red assessed 6 h after LPS or buffer addition. The strong ROS response to LPS exhibited by WT THP-1 cells was blunted in PDCE2 KO cells (n=8, conducted in triplicate on different days). (B) H2O2 release from cells with and without LPS stimulation was assessed in THP-1 WT and KO cells similar to (A) using an Amplex Red/HRP assay to detect extracellular H2O2. The PDCE2 KO cell samples exhibited lower values of extracellular H2O2 than WT cells whether or not they were treated with LPS, and there was no significant response to LPS by the KO cells, unlike WT THP-1 cells (n=12). (C) Cell death was assessed in WT and PDCE2 KO THP-1 cells using Promega Multi-Tox Fluor assay. A significant increase in Dead Cell/Live Cell fluorescence ratio was observed in response to 6 h LPS treatment of WT cells, but this response was dampened in PDCE2 KO cells (n=15). (D) LDH release as a measure of cell death was assessed in WT and PDCE2 KO THP-1 cells using Promega Homogenous Membrane Integrity assay. A significant increase in percent cytotoxicity was observed in response to 18 h LPS treatment of WT cells but not PDCE2 KO cells (n=14). (A-D) All statistical analysis was conducted using one-way ANOVA with Tukey’s multiple comparisons post-test; * p<0.05, ** p<0.01, *** p<0.001.

To compare cellular responses to LPS with or without PDCE2 expression, we used the CRISPR-generated PDCE2 KO THP-1 cells and compared them with the matched WT THP-1 control cells. The KO cells grown in culture exhibited significantly lower levels of PDCE2 protein (Fig. S2), consistent with a mixed population of THP-1 cells of which approximately 60-80% are KO cells. When tested for ROS production responding to LPS treatment, the KO THP-1 cells responded less robustly or not at all compared with WT cells (Fig. 2A and 2B).

Mitochondrial ROS and cell death are linked, thus PDCE2-elicited ROS might play a role in deciding cell fate [20, 21]. Utilizing a commercially available fluorescent live cell/dead cell indicator from Promega, we found that PDCE2 CRISPR KO cells undergo significantly less cell death in response to LPS stimulation than their WT counterparts (Fig. 2C). In addition, measuring cell death by LDH release into the media, we found that PDCE2 KO was protective against cell death caused by LPS (Fig. 2D).

#### 3.2.2 Mutation of PDCE2 Lys residues to prevent lipoylation dampens ROS production and cell death

Covalently bound lipoamide thiol groups of mitochondrial oxo acid dehydrogenase complexes could be sites of S-glutathionylation that block forward or reverse electron flow and upregulate ROS production by PDC [4, 10]. Using our PDCE2 expression system in the CRISPR KO THP-1 cells as above (section 3.1.2 and Fig. 1F), we found that 6 h LPS treatment, which induces PDCE2 glutathionylation (Fig. 1C), also increases ROS production as assessed by MitoSox in all samples (Fig. 3A). However, the LPS-induced increase in ROS detected for WT PDCE2 expressing cells was significantly greater than for the cells transfected with empty vector, whereas the expression of the PDCE2 mutant lacking the lipoylated Lys residues (K132R/K259R) did not enhance ROS production above control (Fig. 3A). Importantly, cell death outcomes as assessed by the Promega live/dead assay closely paralleled ROS production (Fig. 3B).

**Figure 3.**
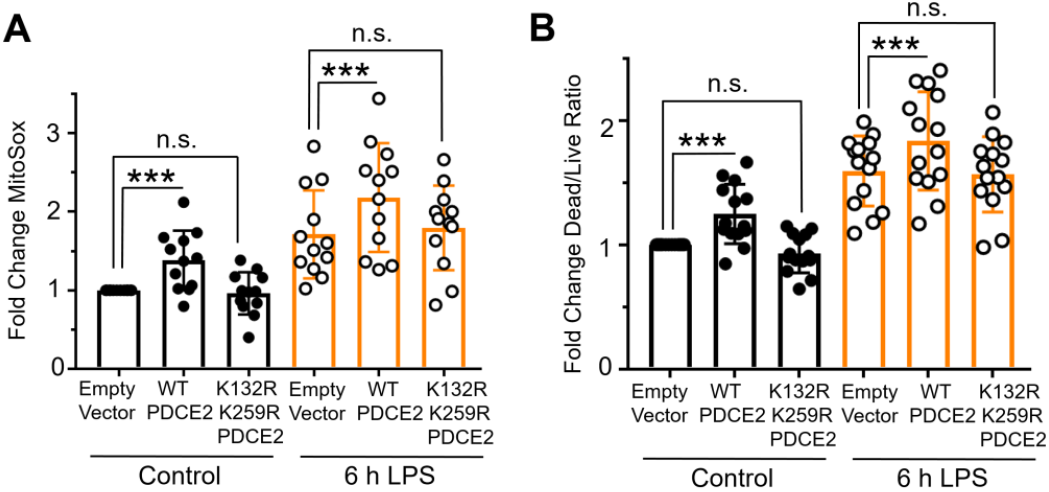
Mutation of PDCE2 Lys residues to prevent lipoylation dampens ROS production and cell death. (A) Mitochondrial ROS production was assessed in PDCE2 KO THP-1 cells transfected with Empty Vector or PDCE2 expression plasmids with and without LPS stimulation using MitoSox Red (n=12, * p<0.05, n.s. = not significant). (B) Cell death was assessed in transfected PDCE2 KO THP-1 cells as in (A), in this case using Promega Multi-Tox Fluor assay (n=14). (A and B) Statistically-significant increases in mitochondrial ROS production and cell death observed upon treatment of WT-expressing cells with LPS were not observed in the cells expressing PDCE2 lacking the lipoyl-linking Lys residues (K132R/K259R). All statistical analysis was conducted using one-way ANOVA with Tukey’s multiple comparisons post-test; * p<0.05, ** p<0.01, *** p<0.001.

### 3.3 Impairment of mitochondrial reductase systems exacerbates LPS-induced PDCE2 glutathionylation, ROS production and cell death, and increases pro-inflammatory cytokine interleukin 1β production, but blunts anti-inflammatory cytokine interleukin 10

#### 3.3.1 MitoCDNB treatment alone does not cause significant PDCE2 glutathionylation but upregulates LPS-induced PDCE2 glutathionylation

Having shown that LPS induces PDCE2 glutathionylation that is associated with greater ROS production and cell death in THP-1 cells, we next searched for ways to biologically induce glutathionylation independent of LPS stimulation to see if alternative mechanisms would lead to the same outcome. To this end, we used mitochondrially-targeted MitoCDNB that compromises both arms of mitochondrial reductase systems (both glutathione-and thioredoxin-based). Formation of glutathione conjugates with the MitoCDNB electrophile in mitochondria catalyzed by glutathione-S-transferase depletes mitochondrial glutathione (GSH), and reactivity of the electrophile with the selenocysteine of thioredoxin reductase-2 in mitochondria compromises the thioredoxin-driven reductase system [22]. Repeating our BioGEE labeling experiments with THP-1 cells, we found that MitoCDNB alone was unable to induce PDCE2 glutathionylation (Fig. 4A). The combined effect of adding both MitoCDNB and LPS together, however, induced a robust increase in PDCE2 glutathionylation that was significantly greater than either alone (Fig 4A).

**Figure 4.**
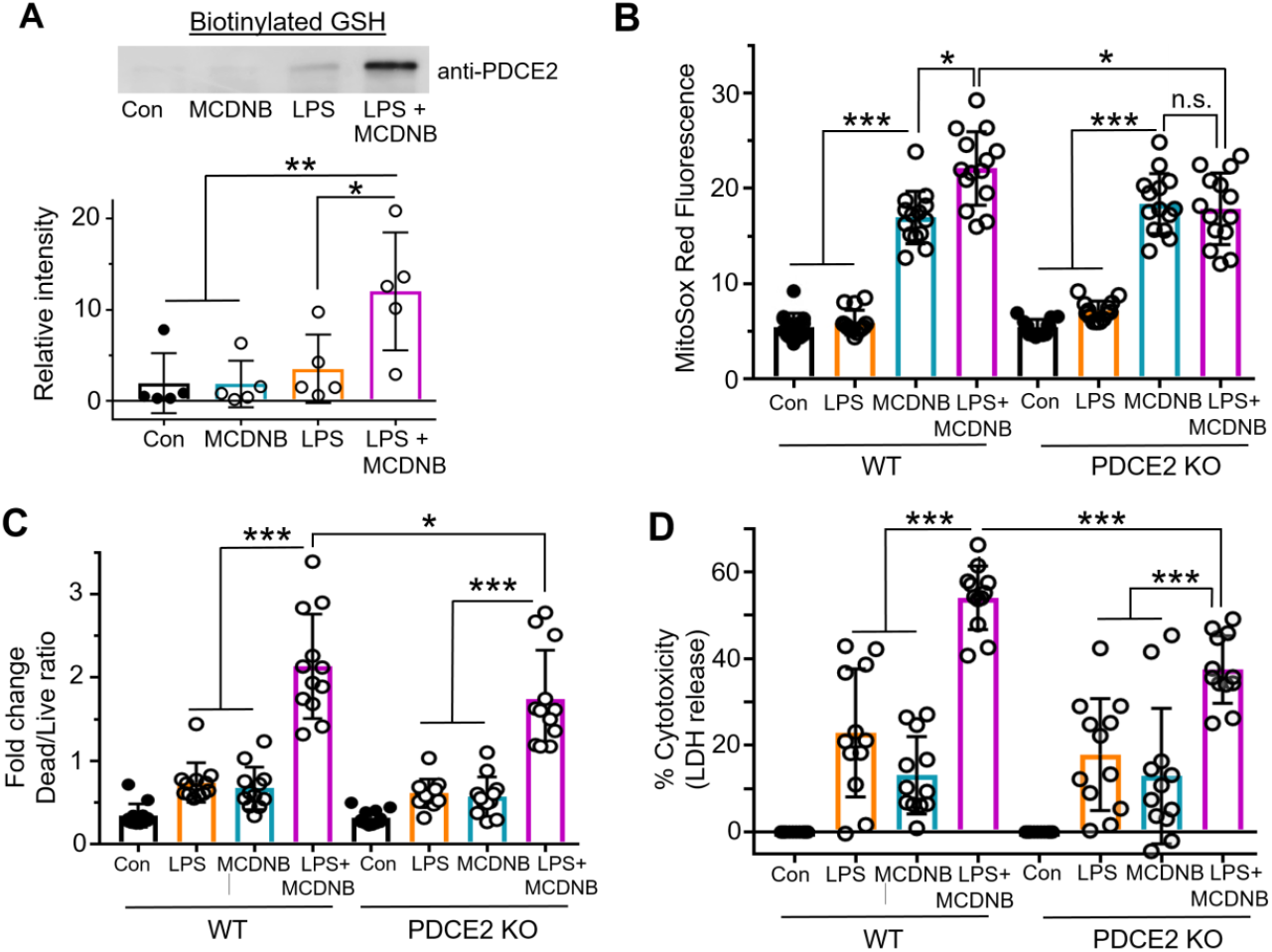
Impairment of mitochondrial reductase systems linked to glutathione and thioredoxin exacerbates LPS-induced PDCE2 glutathionylation, ROS production and cell death. **(A)** BioGEE incorporation assays as conducted in Fig. 1 were used to assess glutathionylation of PDCE2 in THP-1 promonocytes after 6 h treatment with 10 μM MitoCDNB, 1 μg/mL LPS, or a combination of the two (n=5). Using raw densitometry values, a significant increase in glutathionylated PDCE2 was seen with the combined treatments relative to either treatment alone (* p<0.05). **(B)** MitoSox Red was used to assess ROS in WT and PDCE2 KO THP-1 promonocytes. A significant decrease in mitochondrial ROS generation at 4.5 h (shown here) as well as other time points (also shown in Fig. S5) was seen with the combined treatments in PDCE2 CRISPR cells relative to their Wild Type Counterparts (n=14, * p<0.05, ** p<0.01, *** p<0.001). **(C)** Promega Multi-Tox Fluor assay was used to assess cell death in in WT and PDCE2 KO THP-1 cells. The increase in cell death observed with the combined treatments in WT THP-1 cells was significantly less in PDCE2 KO cells (n=12). **(D)** LDH release was assessed as an independent measure of cell death using Promega Homogenous Membrane Integrity assay. Again the LPS-induced increase in LDH release observed with the combined treatments for WT cells was greater that for PDCE2 KO cells (n=12). (A-D) All statistical analysis was conducted using one-way ANOVA with Tukey’s multiple comparisons post-test; * p<0.05, ** p<0.01, *** p<0.001.

#### 3.3.2 MitoCDNB enhances LPS-driven ROS production and Cell Death, but less so in PDCE2 KO THP-1 cells

Evaluating ROS production using MitoSox, we found that MitoCDNB alone induced a significant increase in both WT and PDCE2 CRISPR KO cells (no significant difference between the two), consistent with its mechanism of ROS production being independent of PDCE2. Comparing MitoCDNB-treated cells with and without LPS treatment, ROS levels were significantly enhanced by the LPS treatment in the PDCE2-expressing (WT) cells but were not in the PDCE2 KO cells (Fig. 4B and Fig. S5), suggesting that PDCE2 is required for the LPS-associated effects of the combined treatments. Paralleling the effects seen with ROS production, cell death was significantly blunted in PDCE2 CRISPR KO cells relative to WT cells during combined LPS stimulation and MitoCDNB treatment (Fig. 4C and 4D), suggesting again that PDCE2 is required for the LPS-associated effects of the combined treatments.

#### 3.3.3 Proinflammatory cytokine IL-1β rises, while immunosuppressive IL-10 is strongly suppressed upon co-exposure of THP-1 cells to LPS and MitoCDNB

To further evaluate the link between PDCE2 glutathionylation and inflammatory responses, key cytokines produced during sepsis responses were measured. In particular, pro-inflammatory IL-1β and anti-inflammatory IL-10 were assessed by ELISA from media sampled 6 and 24 hr after the addition of LPS, MitoCDNB, or both. Like ROS levels and cell death, IL-1β produced by LPS-treated THP-1 cells was increased under oxidative stress conditions imposed by MitoCDNB at both 6 and 24 hr (Fig. 5). In sharp contrast, however, anti-inflammatory IL-10 production, which is observed upon LPS treatment in greater amounts relative to MitoCDNB or the control, is completely suppressed with MitoCDNB pretreatment. In every condition, PDCE2 KO cells responded less robustly, but with the same overall pattern as with wild type cells.

**Figure 5.**
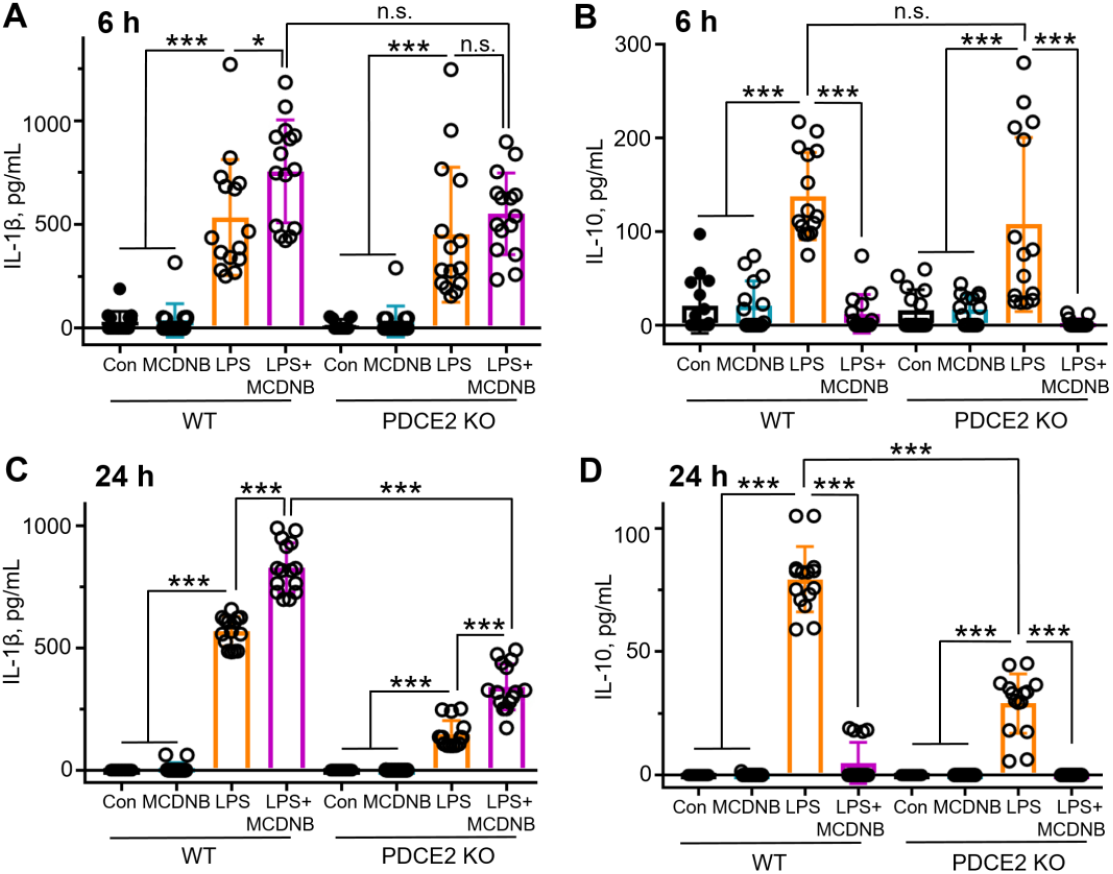
Preincubation of THP-1 cells with MitoCDNB does not induce cytokine production as does LPS addition, but in combination with LPS, leads to a rise in proinflammatory cytokine IL-1β release but strong suppression of anti-inflammatory IL-10 production. **(A-D)** THP-1 cells were treated as in Fig. 4, and media from the cell cultures was sampled at 6 and 24 h after treatments using ELISA kits from BD Biosciences. **(A and C)** Treatment with LPS, but not MitoCDNB, leads to a rise in IL-1β at 6h **(A)** and 24h **(C)**; treatment with both enhances IL-1β production. **(B and D)** Treatment with LPS, but not MitoCDNB, leads to a rise in IL-10 at 6h **(B)** and 24 **(D)**; however, MitoCDNB pretreatment followed by LPS addition leads to little or no IL-10 production in these cells. (n=15 for each panel). (A-D) All statistical analysis was conducted using one-way ANOVA with Tukey’s multiple comparisons post-test; * p<0.05, ** p<0.01, *** p<0.001.

## 4. CONCLUSIONS

Innate immune processes involve multiple complex regulatory networks that intersect to resist pathogens, but which must also be properly regulated, for example in sepsis, where inability to limit tissue damage (disease tolerance) and resolve the inflammatory responses can be fatal. While emerging studies suggest the importance of redox regulatory effects in multiple aspects of these responses, few studies have identified specific redox-related modifications at the molecular level involved in immunomodulation. In tracking a particular oxidative modification, reversible glutathionylation (-SSG formation) occurring within PDC (which serves as a key metabolic node for regulating pyruvate support of the TCA cycle), we found that PDCE2 becomes modified upon LPS stimulation of promonocytes (THP-1 cells in culture) and in isolated human peripheral blood monocytes, likely on the lipoic acid sulfurs that mediate acetyl group transfer in the active enzyme. Thus, the dampening of PDC activity by this redox modification appears to occur in parallel or in addition to the PDCE1α phosphorylation known to downregulate PDC activity during early sepsis. The therapeutic promise of inhibiting this phosphorylation-driven PDC blockage concomitant with normalizing ROS levels as shown recently [23] suggests that mechanisms which help minimize –SSG buildup on PDCE2 could also potentially be harnessed for therapeutic benefit.

Our findings highlight that PDCE2-SSG is generated through LPS signal-mediated, TLR4-dependent processes and not simply enhanced oxidative stress, as the mitochondrial stressor MitoCDNB on its own does not cause PDCE2 glutathionylation. However, when LPS is added to THP-1 cells in which the mitochondrial reductase systems are compromised by MitoCDNB, PDCE2-SSG accumulates to significantly higher levels (likely due to slower reduction, thereby raising steady state levels), an effect which is also associated with higher ROS levels and more cell death (Figs. 4 and 6). This has been established in our cell model of sepsis using chemical probes which give primarily fluorescence readouts of both ROS and cell death; while such measurements cannot be quantitatively converted to concentrations of chemical species or numbers of dying cells, they indicate directions and statistical significance of such changes. Overall, our data suggest that signal-generated PDCE2 glutathionylation is likely an important contributor to adverse outcomes in sepsis.

**Figure 6.**
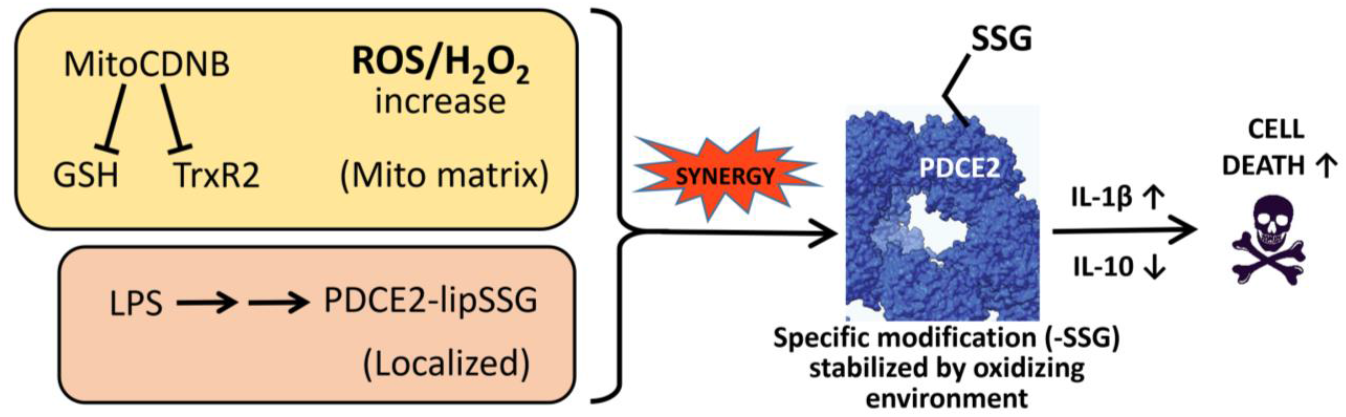
LPS and MitoCDNB synergize to generate enhanced levels of glutathionylated PDCE2, promoting THP-1 cell death through proinflammatory signaling through cytokines.

There is a growing literature linking S-glutathionylation of mitochondrial proteins to the modulation of signaling-relevant ROS production through multiple mechanisms; importantly, release of ROS from cardiac and liver mitochondria is affected by a deficiency in Grx2, a mitochondrial matrix protein which catalyzes deglutathionylation as well as S-glutathionylation of proteins depending on conditions, emphasizing an important role for glutathionylation status [24]. Protein S-glutathionylation was also shown to lower ROS release in skeletal muscle mitochondria through inhibition of pyruvate uptake and modification of complex I [25]. Highly relevant to the present study, S-glutathionylation occurring on PDCE2 of GSSG-treated PDC *in vitro* led to an enhancement of NADH-mediated ROS generation which was reversible by Grx2 treatment, thus demonstrating the link between redox sensing (through protein modification) and ROS production through this complex [10].

This work highlights that redox regulation joins phosphorylation and other modifications (including acetylation) as posttranslational modification mechanisms involved in PDC regulation, and crosstalk between these regulatory layers requires further investigation as well. There is a significant literature showing that oxidative modifications (including glutathionylation) on the active site cysteine of protein tyrosine phosphatases block their activity, elevating phosphorylation levels of their substrate proteins [26, 27], and this may be one regulatory mode in play with PDC regulation. But redox regulation and cross-talk with other modifying enzymes is also much richer and more complex, as multiple systems (including signaling kinases and epigenetic enzymes) are influenced in various ways by their redox status [2, 28-30].

Our findings with respect to immune regulation by cytokines produced by the LPS-stimulated THP-1 promonocytes also have broader implications in health and disease. Importantly, pro-inflammatory cytokines like TNF-α and IL-1β produced by immune cells promote pathogen clearance, but there is also a critical, coordinated production and release of the anti-inflammatory cytokine IL-10 involved in suppression of inflammation and immunopathology, supporting re-establishment of normal function (also referred to as disease tolerance) [31]. Proper balance of inflammation is critical to positive outcomes (survival) in sepsis and many other acute and chronic inflammatory diseases. Our data demonstrated the production of both IL-1β and IL-10 as a result of LPS treatment of the promonocytes, but not when only the mitochondrial stressor MitoCDNB is added. However, the MitoCDNB pretreatment which further exacerbates glutathionylation, ROS production and cell death as a result of LPS signaling also has strong and differential effects on cytokine production. LPS-induced pro-inflammatory signaling is further promoted by MitoCDNB in two ways, by the increase in pro-inflammatory IL-1β, but also by the strong suppression of anti-inflammatory IL-10 (Fig. 5). In the bigger picture, this suggests that compromised reductase systems may be an underappreciated and critical factor in the greater susceptibility and worse outcomes for older or less healthy patients with ongoing redox imbalance due to inflammatory disease or the aging process itself [32]. Thus, the new discoveries in this study move the field of inflammation forward by providing expanded opportunities to understand and even modulate PDC redox status and activity and improve the outcomes of pathological inflammation.

## Supporting information

Compiled Western Blots

## ^1^Abbreviations

Bio-GEE: biotinylated glutathione ethyl ester
DTT: 1,4-dithiothreitol
ETC: electron transport chain
GSH: reduced glutathione
GSSG: oxidized glutathione
LDH: lactate dehydrogenase
LPS: lipopolysaccharide
MitoCDNB: Mitochondrially-targeted 1-chloro-2,4-dinitrobenzene
NEM: N-ethylmaleimide
OGDHC: 2-oxoglutarate dehydrogenase complex
PDC: pyruvate dehydrogenase complex
PDCE2: PDC E2 component
PDCE3: PDC E3 component
ROS: reactive oxygen species
SDS-PAGE: sodium dodecyl sulfate-polyacrylamide gel electrophoresis
TLR: toll-like receptor.

## Author contributions

D.L., L.B.P. and C.E.M. designed the experiments and analyzed the data; experiments were conducted by D.L. under the supervision of L.B.P. and C.E.M. All three authors drafted and edited the manuscript.

## Acknowledgments

The authors thank Drs. Cristina M. Furdui, Co-Director of the Center for Redox Biology and Medicine at Wake Forest University School of Medicine, and Xuewei Zhu who leads the Metabolism and Inflammation Working Group for their input on this project. We also thank Anna Munro for her assistance in gathering data.

## Supplementary Information

Supplementary methods details and data as well as full, uncropped Western blots for this article are provided.

Supplementary Methods and Data includes details of methodology, as well as additional immunoblots and assay results.

Supplementary Compiled Blots includes all Western blots used in the analyses.

