## Supplemental Methods and Data for "Glutathionylation of Pyruvate Dehydrogenase Complex E2 and Inflammatory Cytokine Production During Acute Inflammation Are Magnified By Mitochondrial Oxidative Stress"

### Glutathionylation of Pyruvate Dehydrogenase Complex E2 Protein During Acute Inflammation Is Magnified By Mitochondrial Oxidative Stress, Enhancing Cell Death

David L. Long, Leslie B. Poole*, and Charles E. McCall*

Wake Forest School of Medicine, Winston-Salem, NC 27157 USA

**Supplementary Methods and Data**

**1. Materials and Methods**

**2. Results**

**Figure S1. Pathways for flavin-mediated generation of lipoyl radicals following 1-electron oxidation of fully reduced flavin by molecular oxygen to produce superoxide.**

**Figure S2. Immunoblotting of whole cell lysates to assess PDCE2 expression in the PDCE2 knockout THP-1 population.**

**Figure S3. Confirmation of FLAG-tagged PDCE2 pulldowns for samples immunoblotted for biotin in Fig. 1F.**

**Figure S4. Effect of PDCE2 knockout on LPS-induced $H_2O_2$ production in mitochondria as assessed by MitoPY1.**

**Figure S5. ROS production monitored over time by MitoSox Red fluorescence for WT and PDCE2 KO THP-1 cells.**

#### 1. Materials and Methods

##### Human Promonocytic THP-1 Cell Model

Promonocyte THP-1 cells obtained from the American Type Culture Collection (ATCC) were maintained in complete RPMI 1640 medium (Invitrogen) supplemented with 100 units/ml of penicillin, 100 µg/ml of streptomycin, 2 mM L-glutamine, and 10% fetal bovine serum in a humidified incubator with 5% $CO_2$ at 37 °C. Acute inflammatory responses were induced by stimulating THP-1 cells with 1 µg/mL of Gram-negative bacteria lipopolysaccharide (LPS, Escherichia coli serotype 0111:B4, Sigma Aldrich #L3024) for the indicated times.

##### Human Lymphocyte Studies

Primary lymphocytes were collected from heparinized venous blood samples donated by healthy adult volunteers according to the Institutional Review Board protocol approved by Wake Forest University. Red blood cells, platelets, and polymorphonuclear neutrophils were removed through Isolymph (Gallard-Schlesinger Industries) centrifugation of whole blood. Monocytes were then enriched through a 2 h adherence step, after which nonadherent cells were removed. Cells were then cultured overnight in fresh RPMI 1640 containing 10% FBS prior to stimulation with 100 ng/mL LPS.

##### PDCE2 CRISPR Knockout Cells, Plasmid Constructs and Transfection

PDCE2 CRISPR knockout THP-1 cells were commercially produced by Synthego Corporation and provided along with the matched wild type THP-1 control cell line. The sequence of the guide RNA used for CRISPR was UGGUGCCUGCCUGGCAUUGUG.

To investigate the role of lipoamide thiol groups in redox responses occurring during acute inflammation, a plasmid vector for the expression of PDCE2 in wild type (WT) and mutant forms was designed. PDCE2 has two lysine residues (K132 and K259) where lipoamide cofactors are attached. We designed an N-terminally FLAG-tagged PDCE2 expression plasmid based on SinoBiological plasmid HG15002, synthesized by GenScript, to express PDCE2 WT protein as well as a version where the lysines are mutated to arginines (K132R/K259R), thus leaving the protein incapable of incorporating lipoyl cofactors. Empty vector controls used plasmid CV011 from SinoBiological. Plasmids preparations were conducted using JM109 competent cells (Promega #2005) and the Endotoxin Free Plasmid Purification kit (ThermoFisher #K0861). Cells were transfected with plasmid using GeneX Plus transfection reagent (ATCC #ACS-4004) according to the manufacturer's protocol. In brief, $1x10^6$ THP-1 cells were transfected with 1 µg plasmid that had been preincubated for 15 min with 3 µL GeneX Plus.

##### Detection of Glutathionylated Protein

Cells were preincubated for 1 h with culture medium containing 250 µM BioGEE (ThermoFisher #G36000), a Biotinylated Glutathione analog used to label Glutathionylated proteins, before treatment. For experiments involving mitochondrial glutathione depletion, MitoCDNB (Sigma #SML2573) was added at a final concentration of 10 µM at the same time as BioGEE. Following treatment, cells were washed with ice-cold PBS and lysed in a RIPA buffer (Tris Buffered Saline, pH 7.6, 1% NP-40, 1% Deoxycholate, 0.1% SDS) containing 10 mM *N*-ethylmaleimide (NEM) to block further thiol oxidation. Lysates were passed through a Zeba desalting column (ThermoFisher) to remove unbound NEM, and protein concentration was determined using Pierce 660nm Protein Assay with Ionic Detergent Correction Reagent (ThermoFisher #22660 and #22663). Equivalent amounts of protein from each lysate were incubated for 1 h at 4°C with streptavidin-conjugated magnetic beads (ThermoFisher #21344). Beads were rinsed three times with lysis buffer, and *S*-glutathionylated proteins were released from the beads with boiling for 10 min in Pierce Lane Marker Reducing Sample Buffer (including a final concentration of 20 mM 1,4-dithiothreitol, ThermoFisher #39000). The supernatant was collected from boiled beads, and SDS-PAGE immunoblotting was performed with PDCE2 antibody (GeneTex #GTX109766). In some samples, more stringent washes were also conducted prior to reductive elution to remove less tightly bound proteins associated with the biotinylated (labeled) proteins (1 M NaCl twice, 2 M urea twice, and PBS containing 0.1% SDS); importantly, use of the stringent washes did not change the experimental outcomes.

For experiments involving plasmids, cells were transfected and 36 h later incubated with BioGEE as above. Following LPS stimulation, cells were lysed with IP Lysis buffer (ThermoFisher #87787), and equivalent amounts of protein from each sample were immunoprecipitated with anti-FLAG magnetic beads (ThermoFisher #A36797) for 1 h. FLAG immunoprecipitates were washed three times with IP lysis buffer and proteins were released from washed beads by 10 min incubation with low pH elution buffer (ThermoFisher #21028). Eluates were incubated with an appropriate amount of 5X Pierce Lane Marker NonReducing Sample Buffer (ThermoFisher #39001) for 5 min at 100 °C. SDS-PAGE then immunoblotting was performed with HRP-labeled anti-Biotin antibody (Cell Signaling #7075).

**ROS Assays**

*MitoSox Red.* For endpoint assays, following treatment with LPS and/or MitoCDNB, cells were collected and counted to obtain $2x10^6$ cells per sample, then washed once with Phenol Red Free media and incubated in 5 µM MitoSox Red (ThermoFisher #M36008) for 20 min at 37 °C. Cells were washed after staining, resuspended in 1 mL Phenol Red Free media, then plated in a black 96-well plate, and fluorescence was measured with excitation and emission wavelengths of 510 nm and 580 nm, respectively. Unstained cells were used as a background fluorescence control.

For kinetic measurement of MitoSox Red oxidation during treatments, cells were resuspended in Phenol Red Free media supplemented with 5 µM MitoSox Red at $1x10^6$/mL and incubated 20 min at 37 °C. Cells were spun down and resuspended in Phenol Red Free Media +/- 10 µM MitoCDNB and incubated 30 min at 37 °C. After pretreatment, cells were stimulated +/- LPS and plated in quadruplicate aliquots of 200 µL each in a black 96 well plate. Fluorescence was measured at 30 min intervals in a 37 °C incubator plate reader for 6 h.

*Amplex Red.* A 10 mM stock solution of Amplex Red reagent (Cayman Chemical #10010469) was diluted 1:100 in Phenol Red Free Media. A 1 unit/µL solution of Horseradish Peroxidase (Millipore #516531) was diluted 1:250 into the Amplex Red solution to create the final staining solution. Cells were collected and resuspended in staining solution at $1x10^6$ cells/mL and plated in a black 96-well plate at $2x10^5$ cells per well. Cells were stimulated with LPS, and 6 h later absorbance was measured at 560 nm. A background control of staining solution only was subtracted from each.

*MitoPY1.* Cells at $1x10^6$/mL density were preincubated for 90 min in a 50 µM MitoPY1 (Tocris #4428) staining solution in sterile PBS supplemented with 1 mM Sodium Pyruvate and 5.5 mM Glucose at 37 °C. Following incubation, cells were spun down, resuspended in Phenol Red Free Media, and plated in a black 96-well plate at $2x10^5$ cells per well. Cells were stimulated with LPS, and 6 h later, fluorescence was measured with excitation/emission of $\lambda_{ex}$=488nm/$\lambda_{em}$=544nm. Unstained cells were used as a background fluorescence control.

**Cell Death Assays**

*Fluorescent Live Cell/Dead Cell Assay.* Multiplex Fluorometric Cytotoxicity Assay (Promega #G9200) was used to determine the relative proportion of cell death in the population. Following treatment, triplicate aliquots of cultured cells of 20 µL each were taken from each treatment group and replated into a black 96 well plate. One hundred µL of assay reagent that measures live and dead cells simultaneously was added and plate was incubated at 37 °C for 45 min. Triplicate aliquots of assay reagent only were plated to serve as a background fluorescence control. Following incubation, fluorescence was measured at excitation/emission of $\lambda_{ex}$=400nm/$\lambda_{em}$=505nm for live cells and $\lambda_{ex}$=485nm/$\lambda_{em}$=520nm for dead cells. A ratio of live cell fluorescence to dead cell fluorescence was generated from raw numbers.

*LDH Release Assay.* Homogenous Membrane Integrity Assay (Promega #G7890) was used to measure Lactate Dehydrogenase (LDH) released from cells as a measure of viability according to the manufacturer's instructions. Briefly, aliquots of 5 x $10^5$ cells were plated in individual wells of black 96 well plates and stimulated 18 h with LPS. Plates were equilibrated to room temperature and one row of cells from each treatment group were lysed to generate a max LDH activity. All wells were supplemented with Cyto-Tox One Reagent and incubated for 10 min before fluorescence was measured at excitation/emission of at $\lambda_{ex}$=560nm/$\lambda_{em}$=590nm. Percent cytotoxicity values were obtained for each sample using the untreated WT cells as control (0% cytotoxicity) and the lysed cells for each treatment group as the maximum (100% cytotoxicity).

#### 2. Supplementary Results

##### 2.1 Pathways for generation of lipoyl radicals from PDCE3

Upon reaction of reduced flavin with molecular oxygen through one-electron transfer, in addition to superoxide, a relatively stable flavin semiquinone is formed which can undergo further 1-electron chemistry [1], including reaction with either reduced or oxidized lipoamide from PDCE2 (Fig. S1).

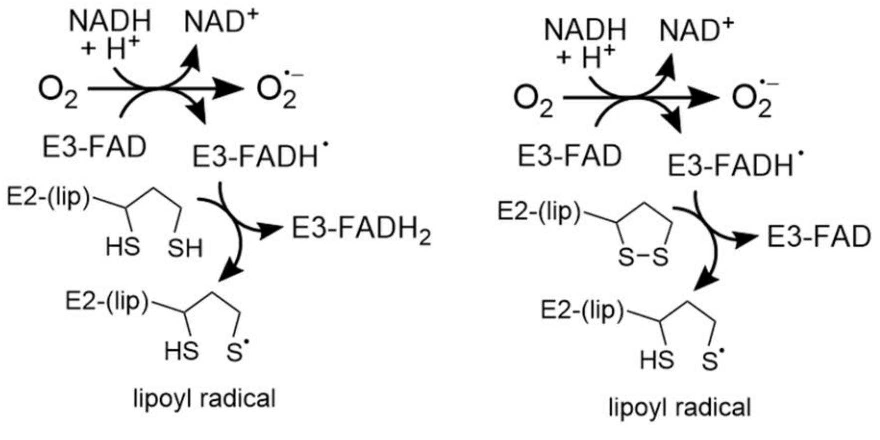


**Figure S1. Pathways for flavin-mediated generation of lipoyl radicals following 1-electron oxidation of fully reduced flavin by molecular oxygen to produce superoxide.**

#### 2.2 Confirmation of PDCE2 knockout (KO), mutant expression and FLAG tag pulldown

CRISPR PDCE2 knockout cells, obtained from Synthego along with the matched WT control, were evaluated for expression of PDCE2 prior to further analysis. We confirmed the strong decrease in PDCE2 protein expression in these THP-1 CRISPR PDCE2 KO cells by immunoblotting lysates collected over a time course of LPS stimulation (**Fig. S2**). Based on the results, PDCE2 KO cells were a significant proportion of the cell population accounting for approximately 60-80% of the total.

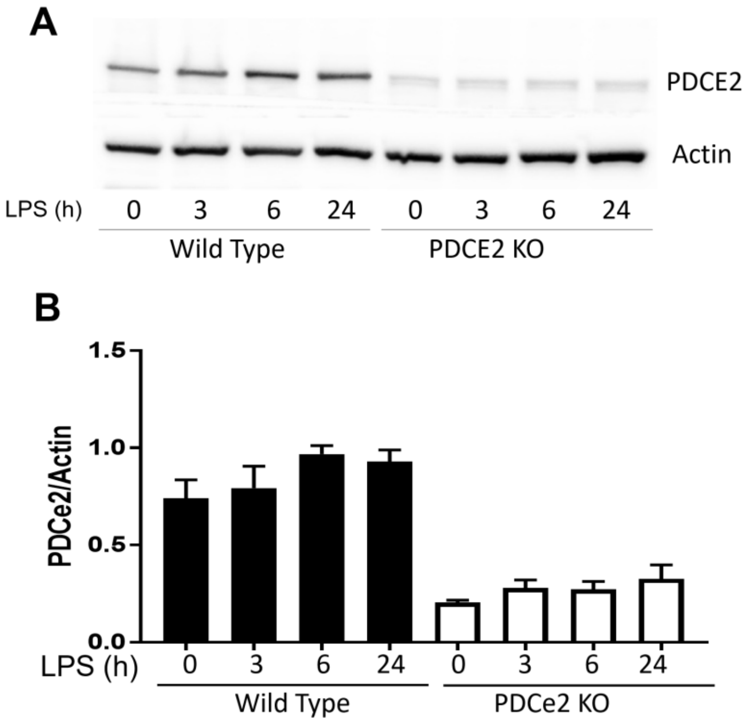


**Figure S2. Immunoblotting of whole cell lysates to assess PDCE2 expression in the PDCE2 knockout THP-1 population.** Wild-Type THP-1 cells and PDCE2 CRISPR THP-1 cells were lysed and immunoblotted after treatment for various times with 1 µg/mL LPS. PDCE2 protein was significantly decreased in the CRISPR cells (n=4, $p<0.05$) relative to Wild-Type counterparts.

The PDCE2 KO cells were further engineered to express FLAG-tagged versions of either WT PDCE2 or a double mutant version of the protein lacking the two lipoyl-bearing Lys residues (K132R/K259R). Following BioGEE labeling and LPS stimulation of these cells, the FLAG-tagged proteins were immunoprecipitated (as shown in **Fig. S3**) and proteins were blotted for biotin as shown in **Fig. 1F**.

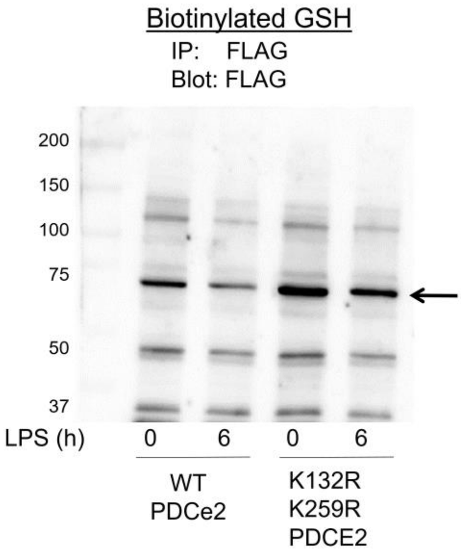


**Figure S3. Confirmation of FLAG-tagged PDCE2 pulldowns for samples immunoblotted for biotin in Fig. 1F.** Following BioGEE labeling and LPS stimulation of WT and K132R/K259R PDCE2 expressing THP-1 cells, immunoprecipitation of the FLAG-tagged proteins was followed by immunoblotting for biotin (shown in Fig. 1F) and for the FLAG tag, shown here (arrow indicates the major band expected for PDCE2).

#### 2.4 Impact of CRISPR-generated knockout of PDCE2 in THP-1 cells on LPS-induced ROS production detected extracellularly by Amplex Red.

Results in **Fig. 2A** and **2B** report the rise in mitochondrial ROS and specifically $H_2O_2$ upon LPS addition to WT PDCE2-expressing THP-1 cells using MitoSox and MitoPY1 fluorescent reporters, respectively. Mutant K132R/K259R PDCE2 expressing THP-1 cells did not exhibit this increase. In support of these results, Amplex Red, which is used to assess extracellular $H_2O_2$, indicated a significant rise in $H_2O_2$ following LPS stimulation for WT THP-1 cells, but not for PDCE2 CRISPR KO cells (Fig. S4).

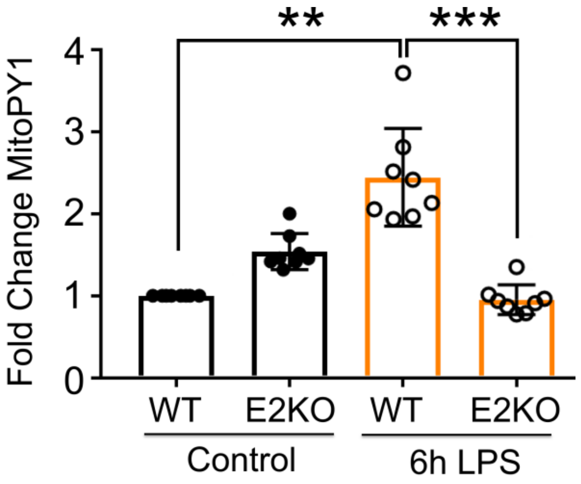


**Figure S4. Effect of PDCE2 knockout on LPS-induced $H_2O_2$ production in mitochondria as assessed by MitoPY1.** The mitochondrially-targeted chemical probe MitoPY1 was used to evaluate mitochondrial $H_2O_2$ production given its selective response to $H_2O_2$ rather than other ROS through release of its boronate protecting group, increasing fluorescence [20]. Following pretreatment with MitoPY1, mitochondrial ROS production was assessed in LPS-treated wild type (WT) THP-1 cells in comparison with CRISPR-mediated PDCE2 knockout (KO) THP-1 cells assessed 6 h after LPS or buffer addition. A significant increase in $H_2O_2$ production was observed in response to 6 h LPS treatment of WT cells, but blunted in PDCE2 KO cells (n=8). Statistical analysis was conducted using one-way ANOVA with Tukey's multiple comparisons post-test; ** p<0.01, *** p<0.001.

##### 2.5 Impairment of mitochondrial reductase systems linked to glutathione and thioredoxin exacerbates LPS-induced PDCE2 ROS production

To track ROS production in LPS and/or MitoCDNB treated cells, cells were pretreated with MitoSox Red, including MitoCDNB for some, then after LPS addition were monitored over 6 hr for fluorescence changes in this fluorescent ROS reporter. Data showed that MitoCDNB alone induced a similar increase in ROS in both WT and PDCE2 CRISPR KO cells (Fig. 4B and Fig. S5) which was higher than that observed with LPS alone. With the addition of both MitoCDNB and LPS, however, the ROS generation was significantly augmented (above MitoCDNB alone) for WT THP-1 cells, but not PDCE2 KO cells, suggesting that PDCE2 is required for the LPS-associated effects of the combined treatments.

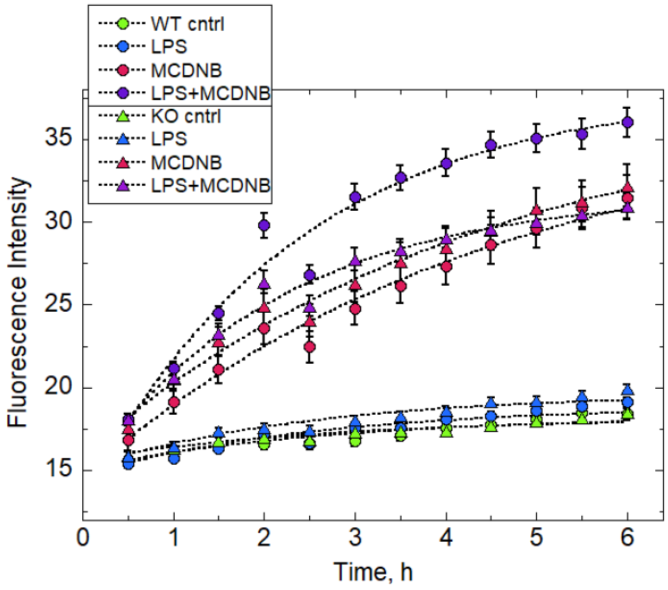


**Figure S5. ROS production monitored over time by MitoSox Red fluorescence for WT and PDCE2 KO THP-1 cells.** Cells pretreated with MitoSox Red and where noted MitoCDNB (MCDNB) were then treated with LPS or buffer and monitored over time in a 37 °C incubator plate reader. Data were fit to a single exponential as a model for processes involved in ROS production after treatments as described by others [2] using Kaleidagraph (Synergy Software).
