## Supplementary material for "Glutathionylation of Pyruvate Dehydrogenase Complex E2 and Inflammatory Cytokine Production During Acute Inflammation Are Magnified By Mitochondrial Oxidative Stress": Compiled Western Blots

### Glutathionylation of Pyruvate Dehydrogenase Complex E2 Protein During Acute Inflammation Is Magnified By Mitochondrial Oxidative Stress, Enhancing Cell Death

David L. Long, Leslie B. Poole*, and Charles E. McCall*

Wake Forest School of Medicine, Winston-Salem, NC 27157 USA

**Supplementary Data showing all full-sized immunoblots**

**Figure S6. Blots for Fig. 1C, THP-1 cells with Bio-GEE labeling and capture, PDCE2 immunoblots.**

**Figure S7. Blots for Fig. 1D, Primary monocytes, Bio-GEE labeling and capture, PDCE2 immunoblots.**

**Figure S8. Blots for Fig. 1F, THP-1 cells with FLAG-tagged PDCE2 expression, Bio-GEE labeling, blot for biotin and for FLAG-tagged PDCE2.**

**Figure S9. Blots for Fig. 4A, MCDNB Bio-GEE labeling and capture, PDCE2 immunoblot.**

**Figure S10. Control blots showing PDCE2 presence in untreated and LPS treated WT and KO cell populations, including β-actin control.**

**Figure S11. Control blots showing FLAG IP proteins after expression of FLAG-tagged PDCE2.**

Fig. S6. Blots for Fig. 1C, THP-1 cells with Bio-GEE labeling and capture, PDCE2 immunoblots

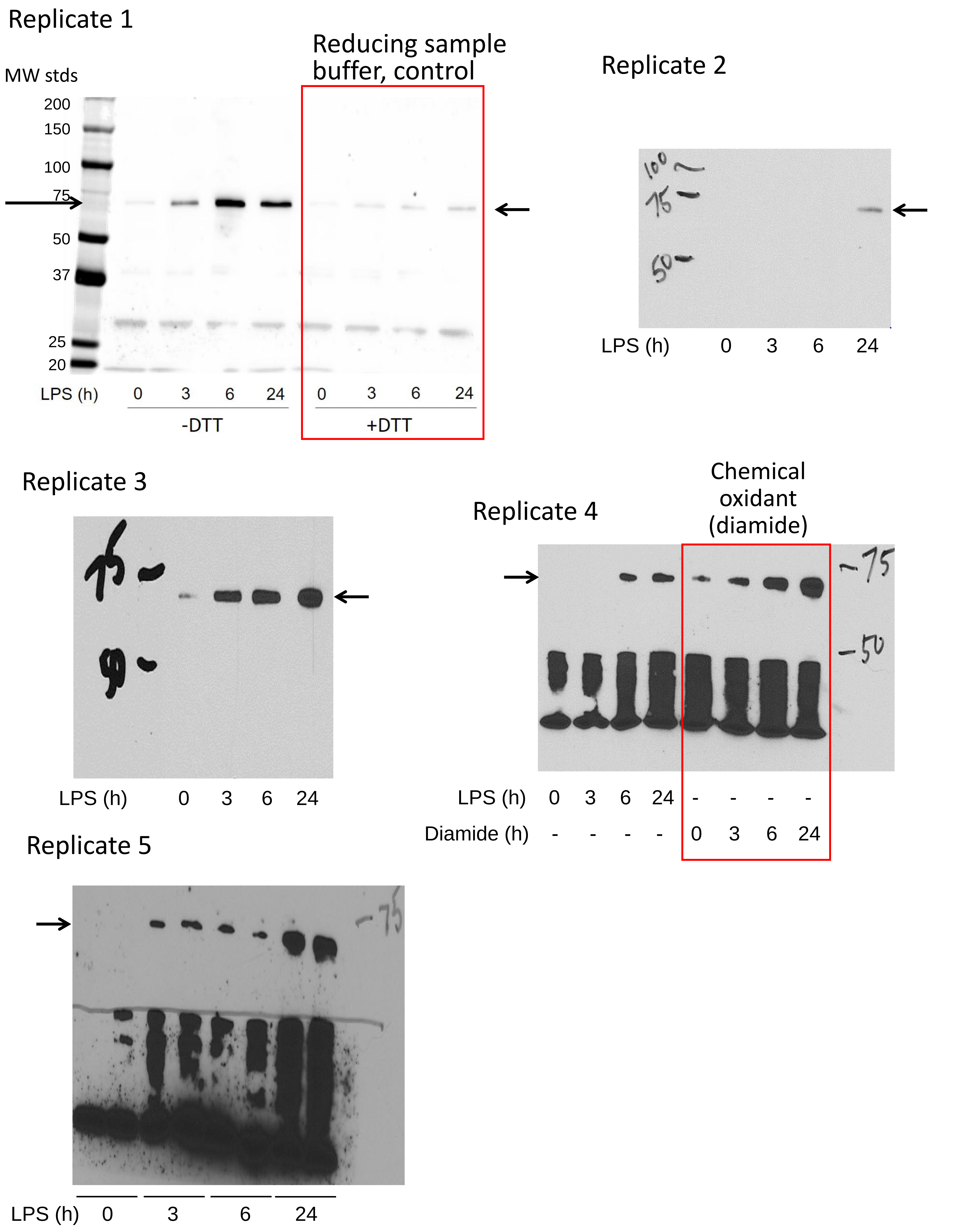

Fig. S7. Blots for Fig. 1D, Primary monocytes, Bio-GEE labeling and capture, PDCE2 immunoblots.

#### Replicate 1

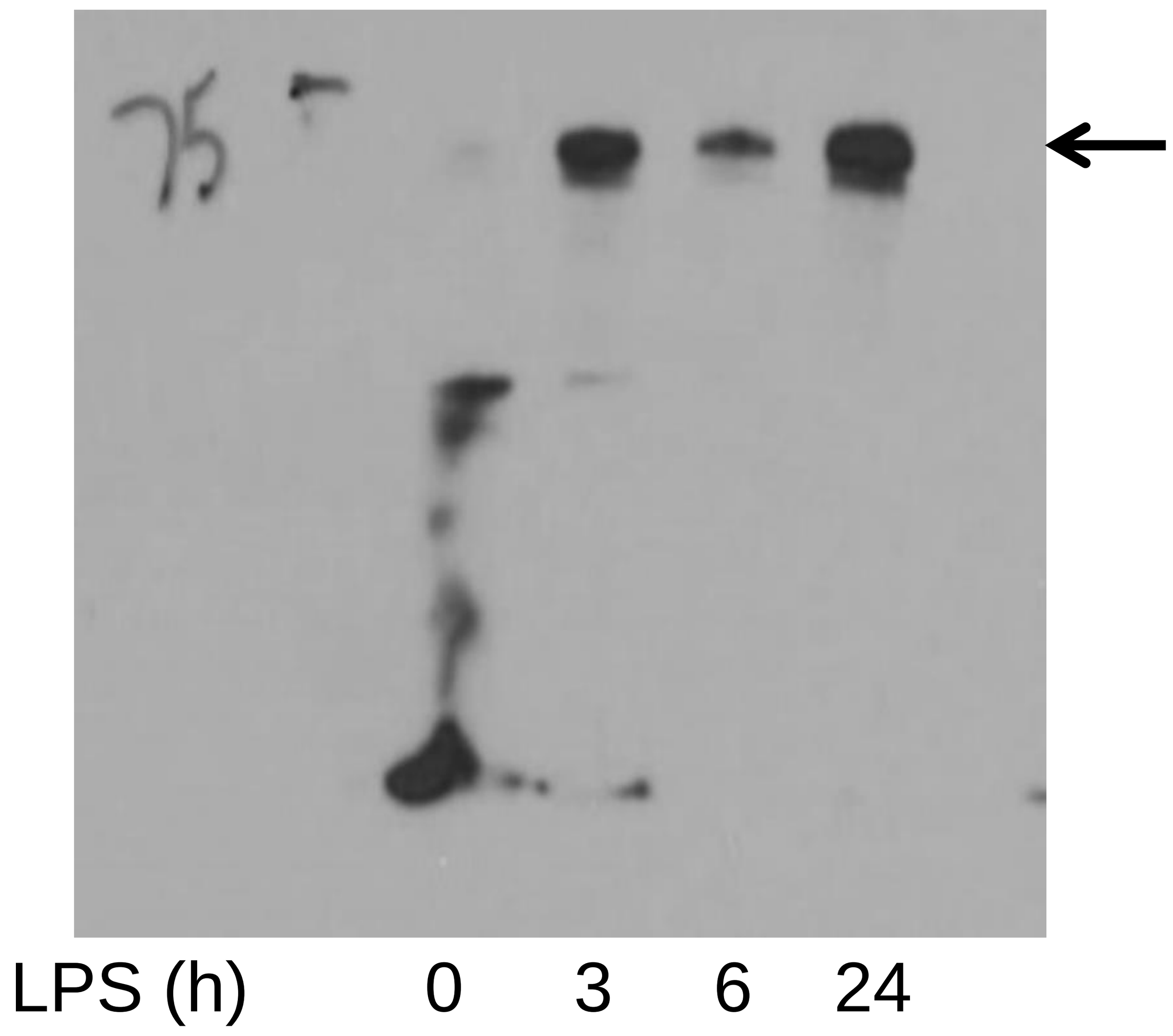


#### Replicate 2

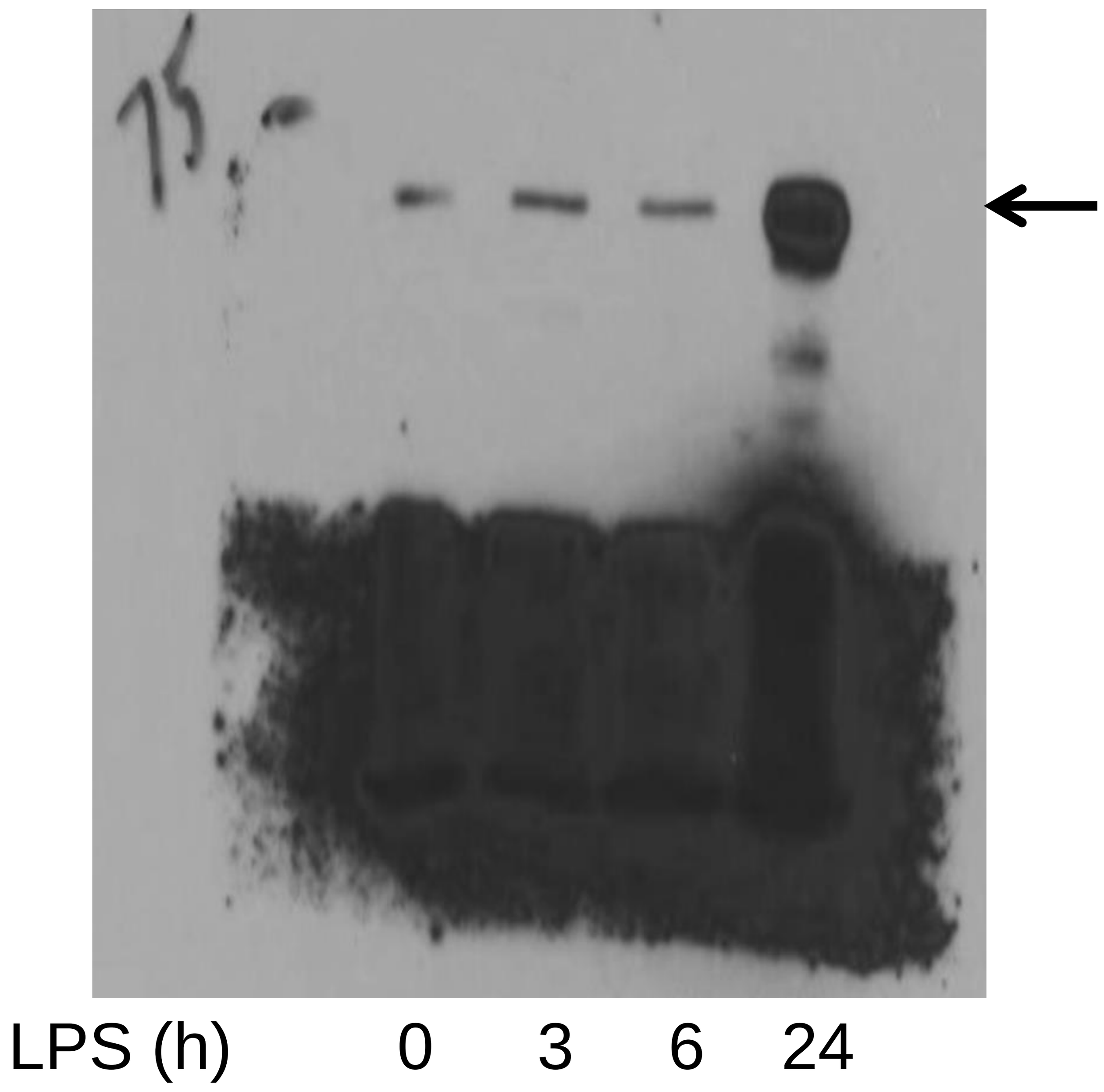


#### Replicate 3

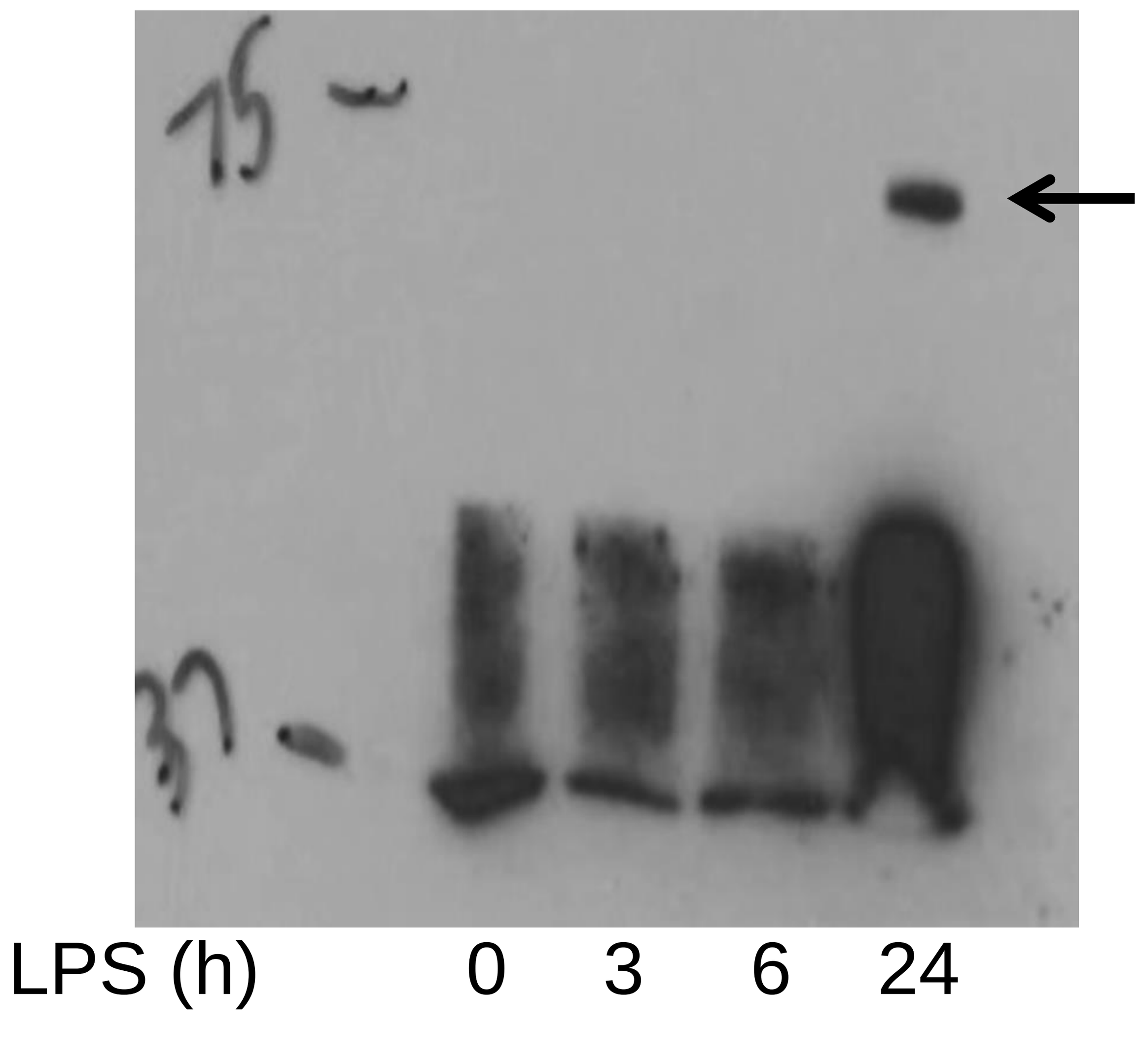

Fig. S8. Blots for Fig. 1F, THP-1 cells with FLAG-tagged PDCE2 expression, Bio-GEE labeling, blot for biotin (left) and blot for FLAG-tagged PDCE2 (right).

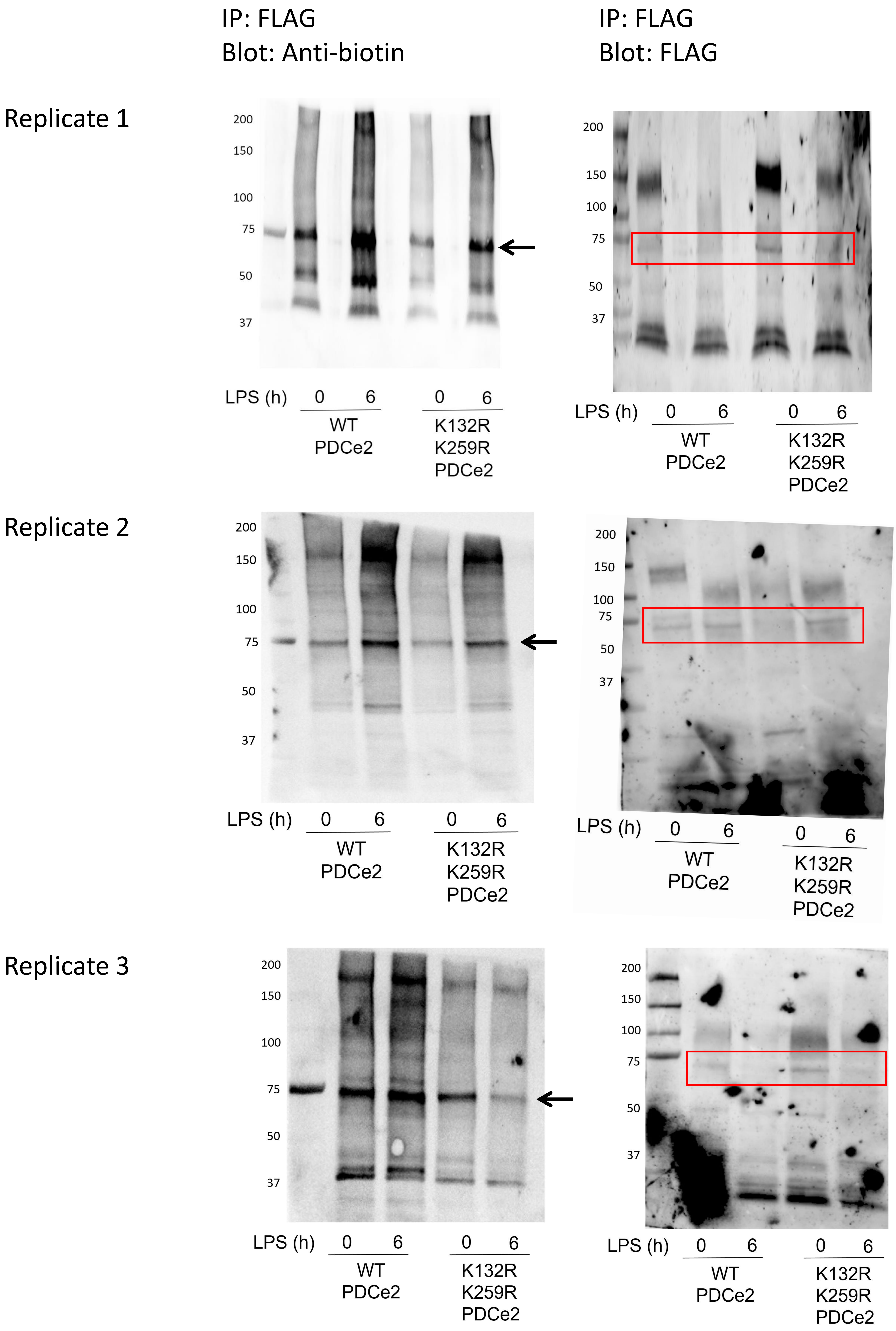

Fig. S8 (continued)

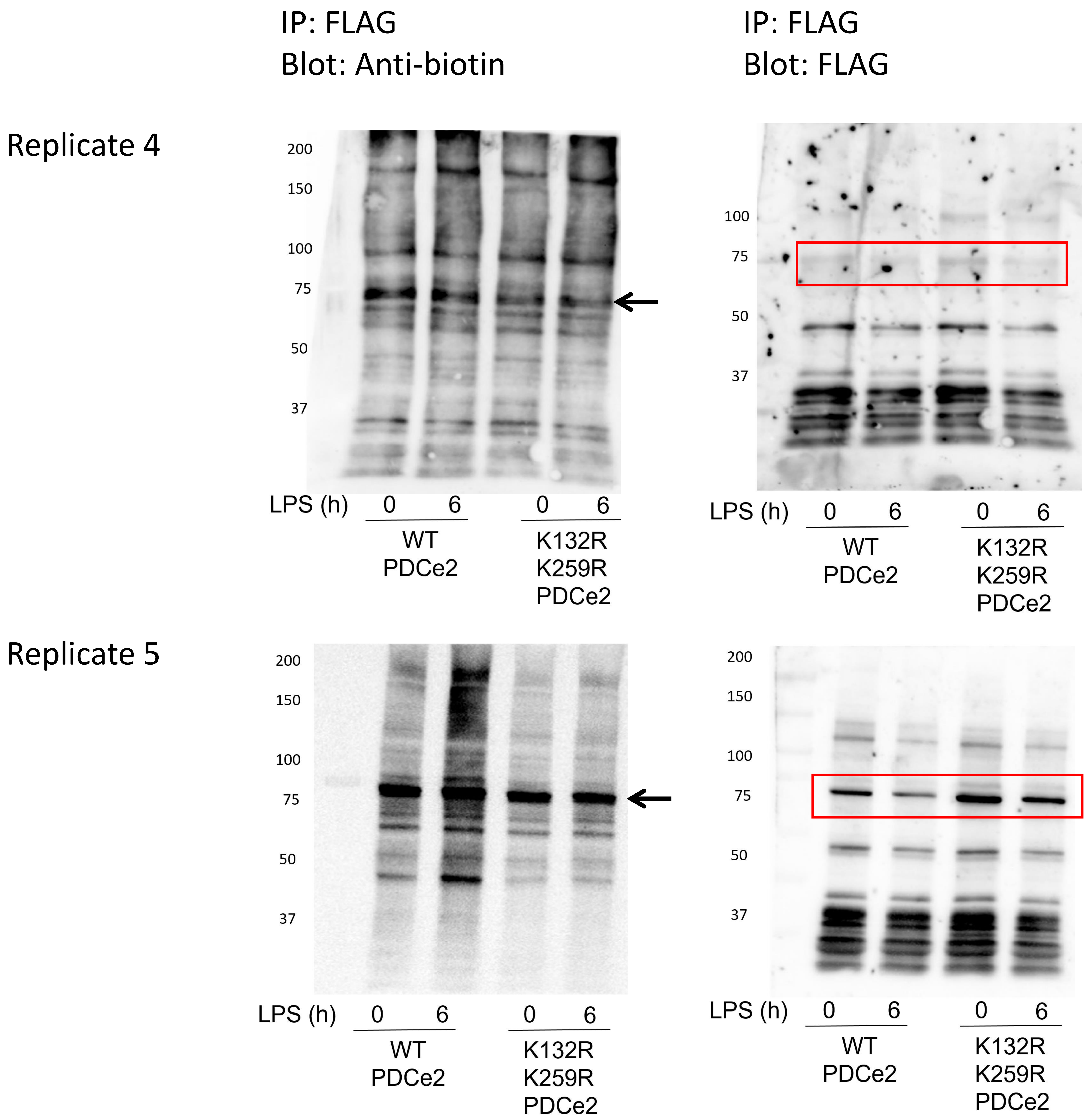

Fig. S9. Blots for Fig. 4A, MCDNB Bio-GEE labeling and capture, PDCE2 immunoblot.

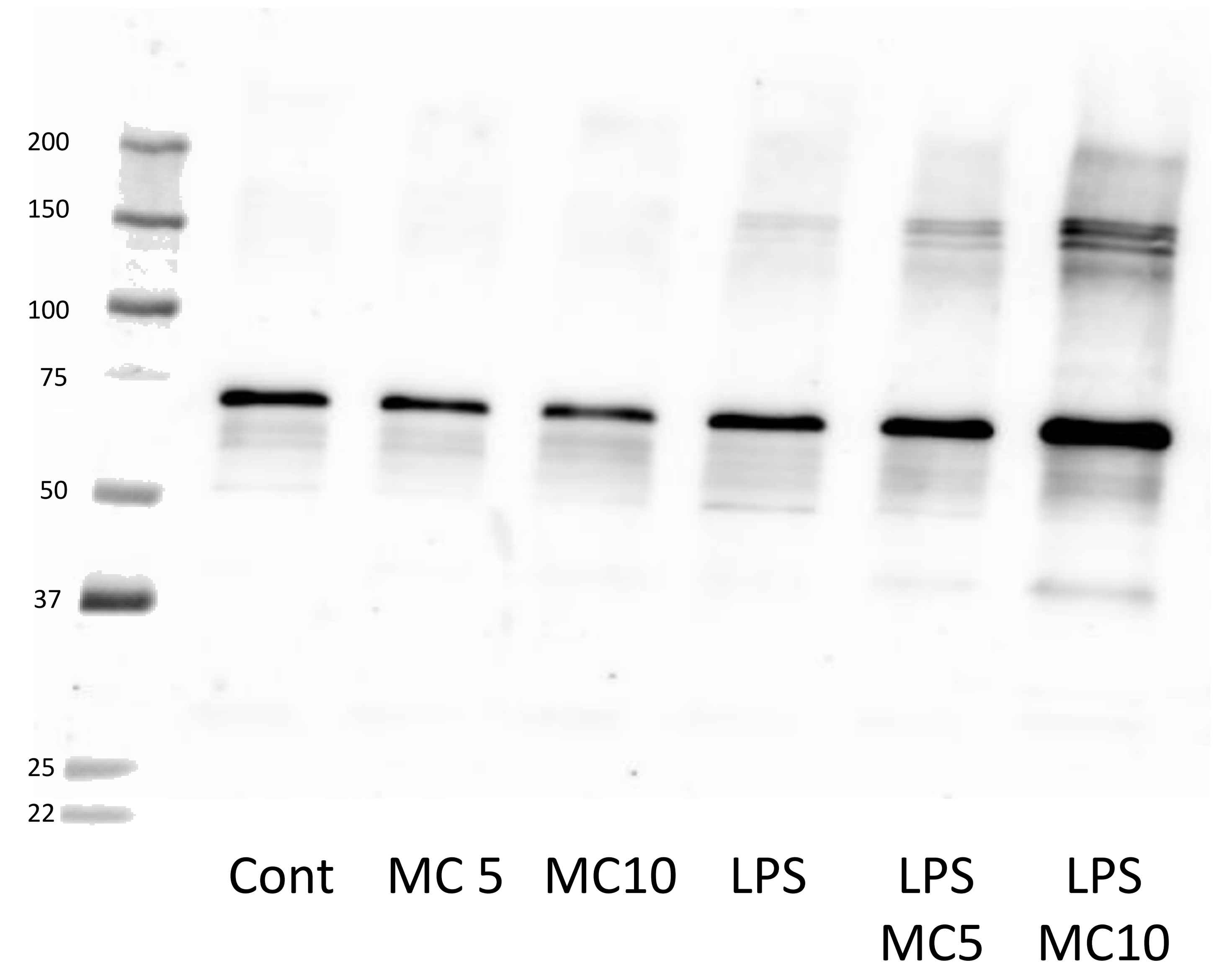


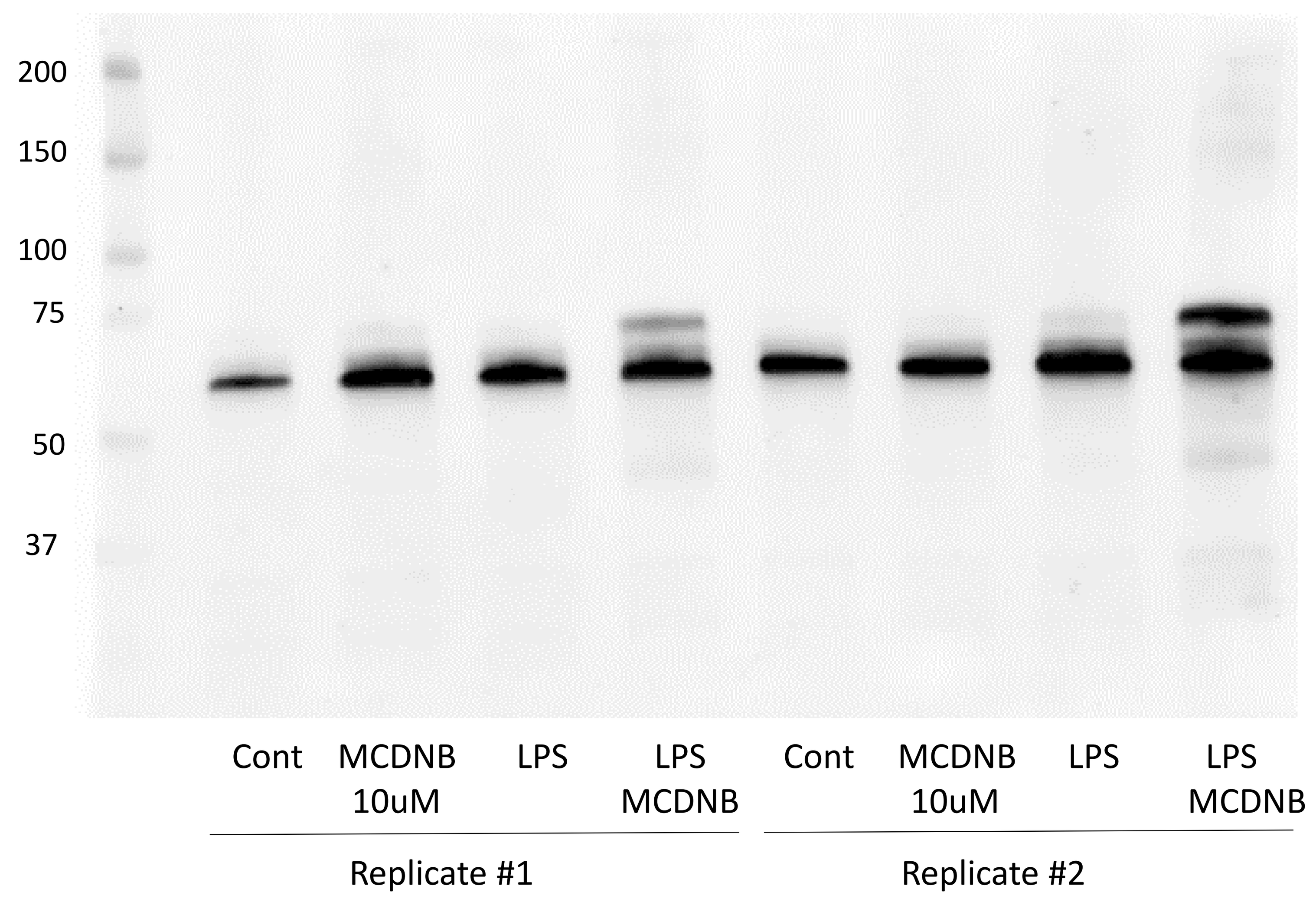


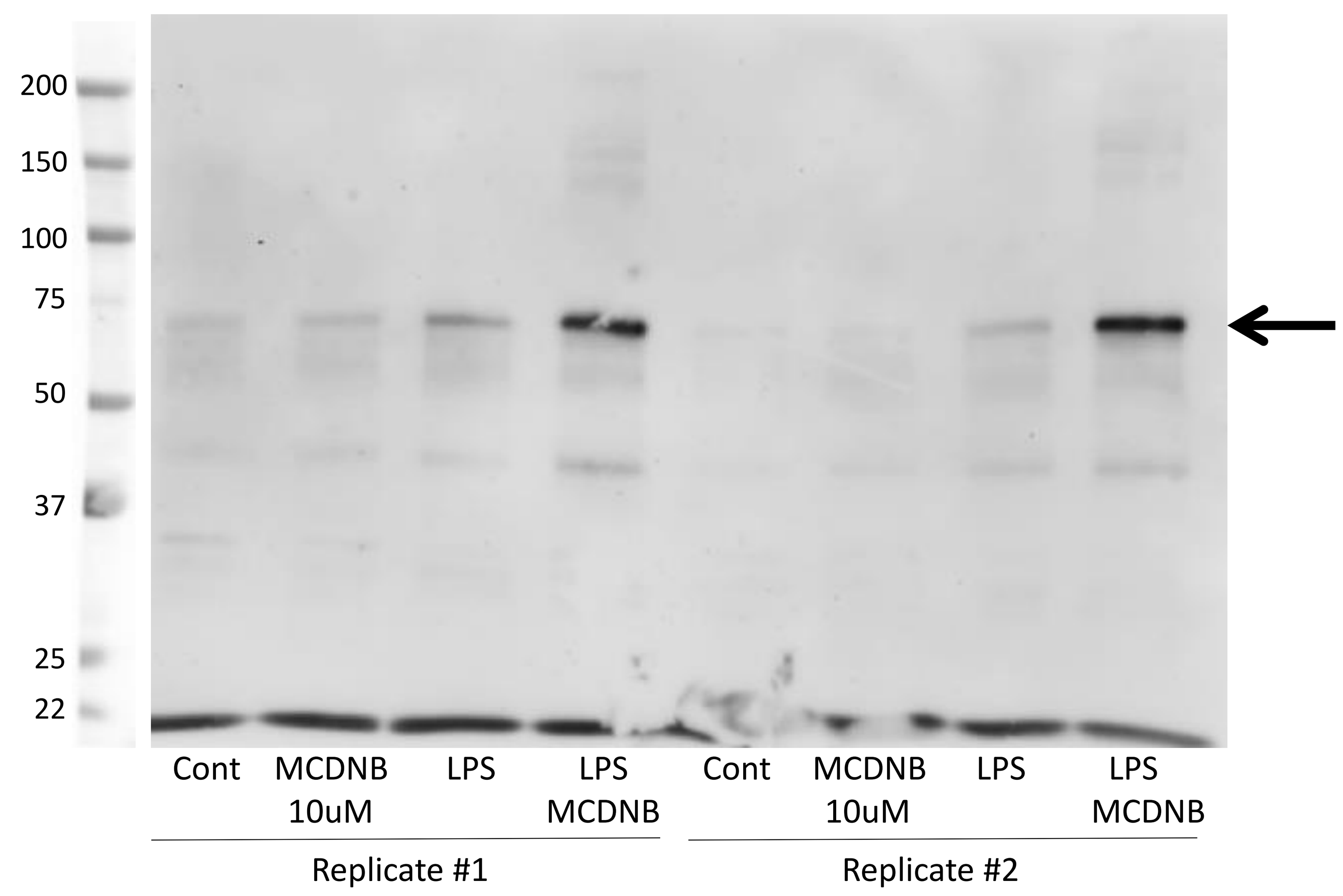

Fig. S10. Control blots showing PDCE2 presence in untreated and LPS treated WT and KO cell populations, including β-actin control.

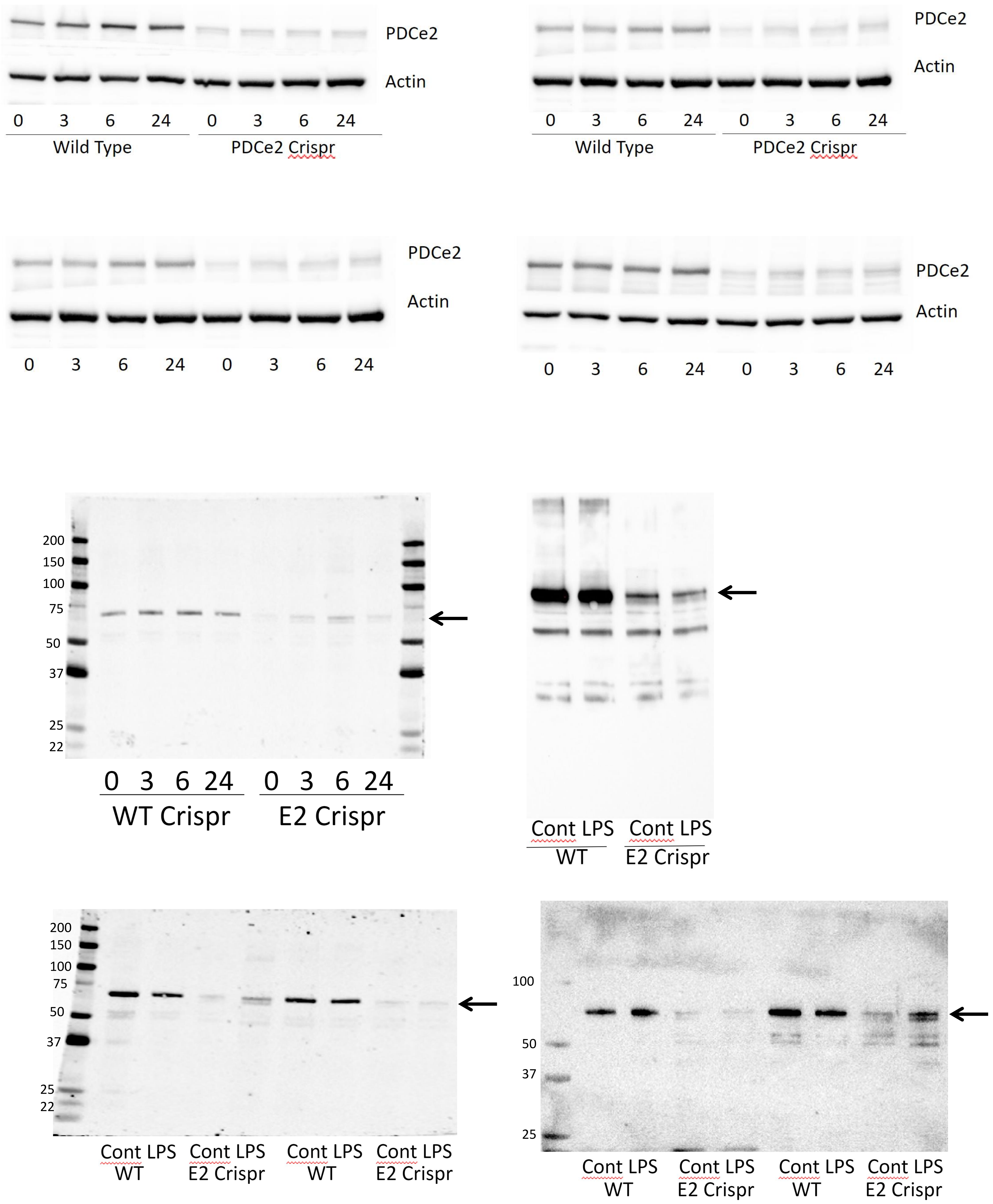

Fig. S11. Control blots showing FLAG IP proteins after expression of FLAG-tagged PDCE2.

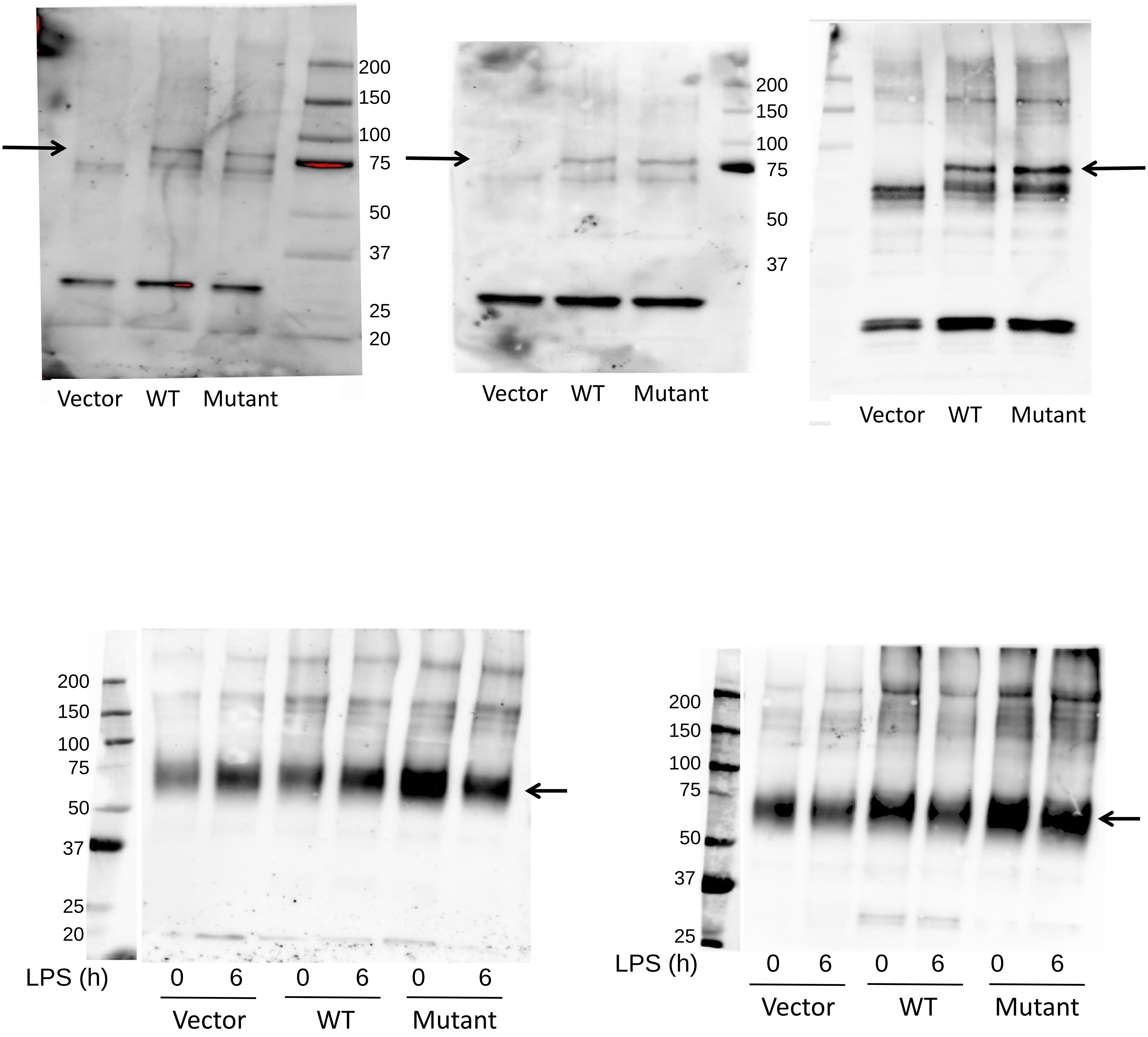
